# Optimizing Lentiviral Vector-Based Delivery of *SCN1A* Transgenes to Mammalian Cells

**DOI:** 10.64898/2026.05.01.722074

**Authors:** Craig Schindewolf, Aguan D. Wei, Franck Kalume, Bruce Torbett

## Abstract

The *SCN1A* gene encodes Na_V_1.1, a voltage-gated sodium channel protein that is necessary for neuronal excitability and whose loss-of-function mutations cause Dravet syndrome, a treatment-resistant childhood onset epilepsy. Gene replacement strategies for this syndrome are challenged by the large size of *SCN1A* and difficulty achieving stable cellular expression. Lentiviral vectors (LVVs) offer sufficient packaging capacity and genomic integration for defective *SCN1A* gene replacement.

Here, we evaluated LVV-mediated delivery of different engineered *SCN1A* transgene sequences in human cells. LVV-transduced cells expressed full-length Na_V_1.1 protein that trafficked to the membrane and produced functional sodium currents. However, *SCN1A* transgene expression declined over time despite stable vector copy number, indicating post-integration regulatory limitations. Expression efficiency varied by *SCN1A* transgene sequence, with a codon-optimized variant showing higher expression despite lower LVV copy number.

Treatment with sodium butyrate, a histone deacetylase inhibitor, significantly enhanced *SCN1A* transgene expression and partially rescued expression decay in a sequence-dependent manner. Incorporation of a ubiquitous chromatin opening element (UCOE) upstream of the promoter to maintain expression resulted in a trend of increased expression and increased responsiveness to butyrate.

These findings demonstrate that sequence-specific and epigenetic factors may influence expression of large transgenes following lentiviral delivery, highlighting key challenges and design considerations for therapeutic *SCN1A* transgene expression.

## Introduction

Voltage-gated sodium channels are essential mediators of electrical signals in the central nervous system^1^. The *SCN1A* gene encodes Na_V_1.1, a large (∼229 kDa) transmembrane protein and sodium channel subunit that is increasingly expressed in the brain^2–5^ during development and plays a critical role in generation of neuronal firing in inhibitory interneurons^6^. Loss-of-function mutations in *SCN1A* are strongly associated with severe epileptic encephalopathies^7^, most notably Dravet syndrome (DS), a developmental and epileptic disorder characterized by early-onset seizures, cognitive impairment, and increased mortality^8^. DS is characterized by drug-resistant, life-long seizures^9^ and therefore, restoration of functional *SCN1A* expression represents a sought-after therapeutic goal^10^.

In recent years, several studies have made remarkable advances in developing therapeutic approaches aimed at supplementing *SCN1A*^10^. These studies have used different methods to overcome several technical challenges. A major concern among these is the large size of the *SCN1A* coding sequence (>6 kb). Lentiviral vectors (LVVs) offer several advantages that make them attractive for delivering large and complex transgenes such as *SCN1A*. LVVs have a relatively large packaging capacity^11^, enabling delivery of full-length sodium channel genes along with regulatory elements. Additionally, LVVs integrate into host genomes, allowing for stable, long-term expression of transgenes, even in nondividing cells^12^, have lower immunogenicity compared to other vectors^13^, and have a strong safety record in clinical use^14^. These features have positioned LVVs as promising tools for gene therapy applications requiring durable expression of large coding sequences. However, despite these advantages, transgene expression from LVVs can be subject to epigenetic silencing over time^15^, and large transgenes may suffer from inefficient mRNA expression^16^ or translation^17^.

In this study, we sought to optimize LVV-mediated delivery of *SCN1A* transgenes to mammalian cells. Building on previously engineered *SCN1A* transgene sequences designed to enhance stability in bacterial systems^18^, we evaluated how sequence composition influences expression efficiency and durability following human cell transduction. We further investigated the extent to which epigenetic modulation—through sodium butyrate treatment^19^ or inclusion of a ubiquitous chromatin opening element (UCOE)^20^—could mitigate the observed decline in *SCN1A* transgene expression over time. Our findings highlight the importance of sequence-dependent effects on transgene expression and underscore the need for continued optimization of vector design to achieve stable and therapeutically relevant expression of large ion channel genes.

## Methods

### SCN1A Transgenes

A previously published *SCN1A* gene sequence (which we refer to as “SCN1A”) was the primary sequence used for this study^18^, obtained through Addgene (#162278). As described in that study, NCBI accession NM_001165963 was used as the basis for this construct, with 5 mutations made (G879A, T897C, T900C, C903T, and T4155C) and a β-globin/IgG chimeric intron intervening sequence (IVS) (GenBank accession U47120) inserted immediately downstream of nucleotide 2946. The “SCN1A” sequence was cloned, using NEBuilder HiFi DNA Assembly (New England Biolabs #E2621), into a pRRL^21^-derived lentiviral transfer plasmid^22^ downstream of a constitutive MND promoter^23^ to drive transgene expression and upstream of a P2A sequence followed by emerald green fluorescent protein (GFP) serving as a reporter. To generate the other plasmids described in the study, the “SCN1A” sequence was substituted with another *SCN1A* construct as previously described^24^ (which we refer to as “Opt”), an IVS-free form of *SCN1A* as previously described^18^ which we kindly received from Alfred L. George, Jr. (which we call “Intron-less”), luciferase, a FLAG-tagged version of “SCN1A”, or *SCN5A* (from Addgene #145374). To generate UCOE-MND-SCN1A-P2A-emGFP, an ubiquitous chromatin opening element of the A2 type^20^ (sequence: ccggaaacacccgaatcaacttctagtcaaattattgttcacgccgcaatgacccacccctggcccgcgtctgtggaactgacccctggtgtacaggagagttcgctgctgaaagtggtcccaaaggggtactagtttttaagctcccaactccccctcccccagcgtctggaggattccacaccctcgcaccgcaggggcgaggaagtgggcggagtccggttttggcgccagccgctgaggctgccaagcagaaaagccaccgctgaggagactccggtcactgtcctcgccccgcctcccccttccctccccttggggaccaccgggcgccacgccgcgaacgcgaactgccgcggtccgcgccgcctccgccctcccccttgggccccaattcccagcgggcgcggcgcgcggcccctccccccgccgggcgcgcgcccgctgccccgcccttcgtggccgcccggcgtgggcggtgccacccctccccccggcggccccgcgcgcagctcccggctccctcccccttcggatgtggcttgagctgtaggcgcggagggccggagacgctgcagacccgcgacccggagcagctcggaggcggtgaacgcgctggctttccttctctctagctctcgctcgctggtggtgcttcagatgccacacgcgt) was cloned immediately upstream of the MND promoter in our original “SCN1A” construct. All of the *SCN1A* constructs used in this study are provided in Sup. Table S1.

### Lentiviral Vector Packaging

Third generation VSV-G-pseudotyped lentiviral vectors were prepared as described previously^25^ with the following modifications: HEK 293T cells maintained in DMEM (Thermo Fisher #11960044) supplemented with 1 x GlutaMax (Thermo Fisher #35050061) with 5% fetal bovine serum (FBS, Omega Scientific #FB-11) were seeded into T182 flasks (Genesee Scientific #25-211) and grown until ∼90% confluent. Cells were then transfected with a mixture of pMDLg/RRE (15 μg), pRSV-Rev (7.6 μg), pMD.2 (VSV-G, 6 μg) and the respective 12.8 – 14.2 kbp transgene plasmid (∼30 – 60 μg) using Profection Mammalian Transfection System (Promega # E1200). Sixteen hours post-transfection, the medium was replaced with the same culture medium containing a lower percentage of FBS (3%). The supernatant was collected two days post-transfection, spun for 5 minutes at 500 *g* to clarify, and ultracentrifuged at 67,865 x *g* for 2 hours and 20 minutes at 4°C on a sterile-filtered 20% sucrose (w/v in Hank’s Balanced Salt Solution) cushion. The pellet was resuspended in Iscove’s Modified Dulbecco’s Medium to a ∼150-fold increase in concentration, aliquoted, and stored at -80°C.

### Western Blot

Cells aspirated of media were lysed in RIPA (Thermo Fisher #89900) with 1:100 protease inhibitor cocktail (Cell Signaling # 5871) and 1:500 benzonase (Millipore Sigma #70664-3) and incubated on ice for 5 minutes. Lysates were then sheared by repeated aspirations (∼4 times) through a 23g needle, spun down at max speed at 4C in a microcentrifuge for 10 min, and the liquid portions combined 1:1 with fresh 2x Laemmli Sample Buffer (Bio-Rad #1610737) containing 5% 2-Mercaptoethanol (Bio-Rad #1610710). Samples were incubated at 95C for 5 minutes before running standard SDS-PAGE. After electrophoresis, the gel was incubated in transfer buffer (1x Tris-glycine [3g/14.4g respectively per L)] containing 15% ethanol and 0.0075% SDS) and transferred to ethanol-activated PVDF membrane overnight at constant 10V at 4C. The membrane was then blocked for ∼30 minutes with a 5% blocker (Bio-Rad #1706404) solution in TBS-T, washed briefly with TBS-T, then incubated with primary antibody (either Sigma #F1804 [anti-FLAG] at 1:500 dilution, Antibodies Incorporated #75-023 [anti-Na_V_1.1] at 1:500 dilution, or Thermo Fisher #MA1-744 [anti-actin] at 1:2500 dilution) overnight at 4C. After, membranes were washed 3x with TBS-T for 5 minutes each wash and incubated with anti-mouse HRP-conjugated secondary antibody (Jackson Immunoresearch #115-035-003) at 1:3000 dilution. After another 3 washes in TBS-T for 5 minutes each wash, membranes were developed with ClarityMax (Bio-Rad #1705062).

### Immunocytochemistry

Cells aspirated of media were fixed with BD Cytofix/Cytoperm (BD Biosciences #554722) for ∼30 minutes and then blocked with 10% blocker (Bio-Rad #1706404) in PBS for ∼1 hour. Blocker was aspirated and cells were then incubated overnight at 4C with 1:138 anti-Na_V_1.1-FL550 conjugate (Antibodies Incorporated #75-023-FL550), diluted in 2% milk in PBS. The next day, cells were washed 3 x with 1% milk in PBS, covered with mountant (Thermo Fisher #S36968), overlaid with glass cover slips, and imaged via microscopy.

### Flow Cytometry

Single cell suspensions in DPBS from cultures were used for flow cytometry, particularly amount of fluorescent signal at 495 – 535 nm upon 488 nm excitation. Cells were analyzed on a FACSymphony A5 (BD Biosciences) and then data was analyzed with either FlowJo software or Floreada (https://floreada.io). For samples that received sodium butyrate, 5 mM sodium butyrate (final concentration) was added to cultures ∼24 hours before flow cytometry.

### Relative Vector Copy Number Analysis via Quantitative PCR

Genomic DNA was purified from cells using PureLink (Thermo Fisher #K182001). DNA was amplified using PowerUp SYBR Green Master Mix (Thermo Fisher #A25741) with primers targeting vector *emGFP* (forward: 5’-taaacggccacaagttcagc and reverse: 5’-gtaggtcaaggtggtcacga) and *MKL2* (forward: 5’-agatcagaagggtgagaagaatg and reverse: 5’-ggatggtctggtagttgtagtg). Relative vector copy was computed as the ratios between samples of the difference in threshold cycles of vector target (*emGFP*) and human reference (*MKL2*) (i.e. ΔΔ𝐶𝑡 method).

### Electrophysiology

Previously transduced cells were dissociated with TrypLE (Gibco #12-604-013) and replated at low density onto 12 mm poly-D-lysine coated glass coverslips (NeuVitro# GG-12-pdl) in 24-well tissue culture plates (Fisher Scientific #FB012929) for patch-clamp electrophysiology. Typically, ∼10,000 – 15,000 cells were replated per well at sufficiently low density to isolate individual cells. This was necessary to prevent the formation of electrical junctions between contacting cells, which precludes adequate space-clamp recording conditions. Recordings were performed from 0.5 – 3 days after replating at low densities.

For patch-clamp recordings, coverslips containing adherent transfected cells were transferred to the stage of a Zeiss AxoExaminer.A1 microscope equipped with an 40X water immersion objective and epifluorescence capability. Pipettes were positioned with a Sutter MPC-325 micromanipulator. Whole-cell voltage-clamp recordings were acquired with an AxoClamp200B amplifier (Molecular Devices) using pClamp10.4. The composition of recording solutions was as previously described^26^—bath (in mM): 140 NaCl, 2 CaCl_2_, 2 MgCl_2_, 10 HEPES, pH 7.4); pipette internal solution (in mM): 35 NaCl, 105 CsF, 10 EGTA, 10 HEPES, pH 7.4). Patch pipettes were pulled from borosilicate glass (World Precision Instruments #1B120F-4) on a P-97 Sutter Instruments puller, and fire-polished on a Micro-Forge MF-830 (Narashige International USA) to a resistance of 0.8 – 1.5 MΩ. Currents were allowed 5 – 10 mins to stabilize after achieving whole-cell recording configuration, and acquired at 50 kHz, filtered at 5 kHz. Peak currents were recorded in response to a family of voltage steps from a holding potential of –120 mV to 40 mV, in 5 mV increments, with an inter-pulse interval of 2 seconds to allow channels to fully deactivate to the deep closed state. Capacitive transients were subtracted using a P/4 subtraction scheme, employing a current subtraction template derived from scaling the voltage command protocol by one-fourth. Series resistance compensation was >90% for all recordings. Current traces were analyzed and plotted using pClamp10.4 (Molecular Devices) and Origin 8.5.

## Results

### The full-length *SCN1A* transgene is expressed and traffics to the membrane

Because DNA constructs encoding large ion channel proteins are historically hard to propagate in *E. coli* due to sequence stability, we relied on a previously published *SCN1A* sequence that introduced silent mutations and a strategically placed chimeric intron to disrupt cryptic prokaryotic promoters^18^, as depicted in Fig. 1a (full sequence in Sup. Table S1). We verified that the *SCN1A* sequence expressed full-length Na_V_1.1 protein in a human cell line, HEK 293T, when transiently overexpressed (Fig. 1b). We also observed trafficking to the membrane, consistent with the function of Na_V_1.1 as an ion channel (Fig. 1c).

**Figure 1:**
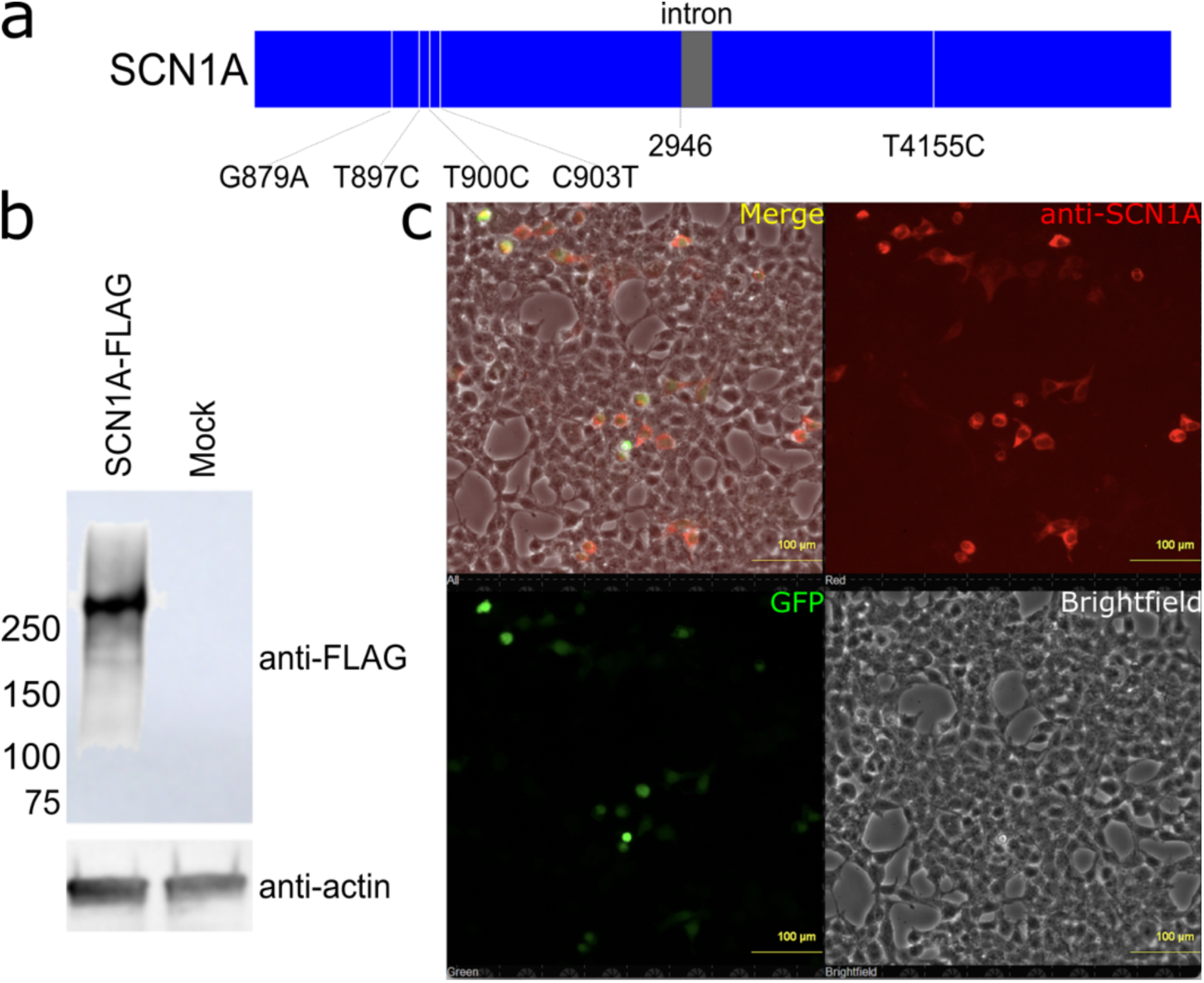
SCN1A is expressed in full length and traffics to the membrane. (a) Schematic of SCN1A open reading frame. NCBI accession NM_001165963 was used with nucleotide mutations and a β-globin/IgG chimeric intron intervening sequence (IVS) (GenBank accession U47120) immediately downstream of nucleotide 2946 denoted as previously described^18^. (b) Western blot of 293T cells transfected with C-terminally FLAG-tagged SCN1A or mock cells, stained with anti-FLAG primary antibody (top) as well as anti-β-actin (bottom, same samples stained separately). A color image containing the protein molecular weight ladder (not shown) was overlaid on the black-and-white chemiluminescent image. (c) 293T cells transfected with a plasmid co-expressing SCN1A and GFP were fixed in 4% paraformaldehyde and stained with anti-Na_V_1.1 conjugated with a 550 nm-emitting fluorophore.

### *SCN1A* transgene expression strength is sequence-dependent and decays over time

Having validated the expression and function of the *SCN1A* construct we were using, we then cloned the sequence into a self-inactivating lentiviral transfer vector^22^, originally derived from pRRL^21^. We included a self-cleaving P2A sequence^27^ and emerald green fluorescent protein (emGFP)^28^ downstream of *SCN1A*, all driven by the constitutive viral-derived *m*yeloproliferative sarcoma virus enhancer, *n*egative control region deleted, *d*l587rev primer-binding site substituted (MND) promoter^23^. To directly compare the performance of different published *SCN1A* transgene sequences, we also cloned an intron-less *SCN1A* construct^18^ (referred to as “Intron-less”), as well as a codon-optimized version of *SCN1A* published in a recent study (referred to as “Opt”)^24^. The “Opt” sequence differs from our originally tested construct (which we refer to simply as “SCN1A”) due to: 1) the sequence and position (proximal to the start codon) of the chimeric intron, 2) a shorter, more widely expressed form of exon 11 that makes the Na_V_1.1 protein 11 amino acids shorter, and 3) a threonine at position 1056 that corresponds to the predominant allele (https://gnomad.broadinstitute.org/variant/2-166036278-C-T and Sup. Table S1). For cellular expression assessment of the different *SCN1A* sequences present in LVVs, all were packaged as described in the Methods and concentrated 150-fold. We also packaged a control LVV encoding luciferase as a coding gene substitute for the *SCN1A* transgene (Fig. 2a and Sup. Table S1). We were able to verify function of the LVV encoding the “SCN1A” sequence in transduced 293T cells, observing current production upon voltage stimulation, consistent with voltage-gated ion current facilitated by channel expression (Fig. 2b).

**Figure 2:**
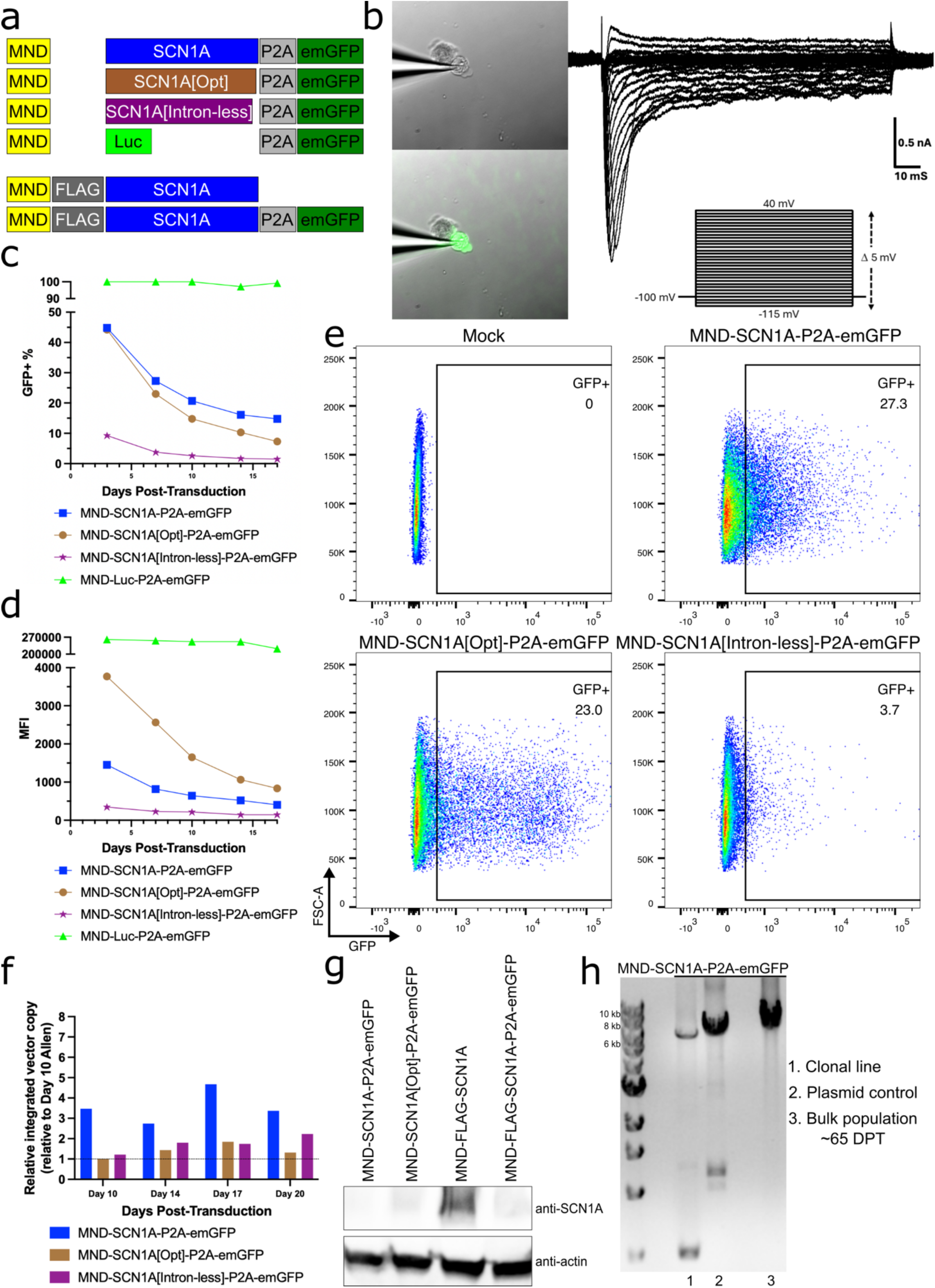
SCN1A construct expression strength is sequence-dependent and decays over time. (a) Sequences of *SCN1A* constructs constructed and packaged into lentiviral vectors and transduced into 293T cells. MND = MND promoter. SCN1A = *SCN1A* sequence as in Figure 1, which we refer to as “SCN1A” in the text. SCN1A[Opt] = construct as previously described^24^. SCN1A[Intron-less] = construct as previously described^18^. Luc = luciferase. (b) Currents evoked in a 293T cell transduced with MND-SCN1A-P2A-emGFP, in response to a family of step depolarizations from a holding potential of - 115V to 40V, in 5-mV increments. (c) Percent of GFP+ and (d) mean fluorescence intensity (MFI) of transduced cells over time. (e) Flow plots at 7 days post-transduction (DPT). (f) Quantification of integrated vector copy, relative to cells transduced with MND-SCN1A[Opt]-P2A-emGFP at 10 DPT (set to 1). (g) Western blot of SCN1A in transduced cells at 8 (first three samples) or 9 (last sample) DPT. (h) Amplification of genomic DNA with primers targeting the MND promoter and emGFP, which spans the entire *SCN1A* transgene sequence. Rightmost lane: bulk 293T cells transduced with MND-SCN1A-P2A-emGFP at 65 days post-transduction. “Plasmid control”: MND-SCN1A-P2A-emGFP lentiviral transfer plasmid itself (i.e. not lentiviral vector-transduced cells) as positive control. “Clonal line”: a subline of the bulk MND-SCN1A-P2A-emGFP-transduced 293T that was particularly emGFP-bright.

Next, we assessed how well LVVs with different *SCN1A* transgene sequences expressed when we transduced 293T cells. For all but the luciferase control LVV, we found that expression of our *SCN1A* transgenes (as measured by emGFP fluorescence) declined over the course of 17 days (Fig. 2c, d). However, this was not due to a decrease in LVV copy (Fig. 2f). The Opt LVV showed the highest mean fluorescent intensity (MFI)(Fig. 2d, e), despite having lower copy number relative to the “SCN1A” and Intron-less LVVs. To verify that transduced cells were expressing the Na_V_1.1 protein (and not just emGFP) from our LVV-transduced cells, we also quantified Na_V_1.1 protein and detected it in both the cells transduced with Opt or versions of “SCN1A” with a FLAG tag, verifying that the LVVs were producing full-length Na_V_1.1 protein (Fig. 2g). Lastly, we assessed LVV sequence integrity in transduced cells at around 65 days post-transduction. For cells transduced with the “SCN1A” LVV, we saw a clean band from PCR amplification with primers targeting the MND promoter and the emGFP reporter, suggesting no intervening truncations, i.e. in the *SCN1A* transgene sequence itself (Fig. 2h, rightmost lane). Interestingly, we did note that an emGFP-bright clonal population derived from these cells showed evidence of a deletion of a large part of the *SCN1A* transgene sequence (Fig. 2h, “Clonal line”) which we sequence verified. This suggests that over time in culture, rearrangement of the transduced *SCN1A* transgene can result in loss of function, but does not appear to be the predominant outcome and does not fully explain the loss of *SCN1A* transgene expression over time.

### Butyrate helps counter the decay of *SCN1A* and *SCN5A* transgene expression in a sequence-dependent fashion

Given we observed that LVV copy number for the different *SCN1A* LVVs did not decline over time, although expression of the emGFP reporter did, we tested whether rescue of *SCN1A* transgene expression would occur from altering epigenetic regulation. Butyrate is a short-chain fatty acid known to inhibit histone deacetylation, which is proposed to promote gene expression^19^. Addition of 5 mM sodium butyrate countered the expression decay of the emGFP reporter signal in *SCN1A*-transgene LVV-transduced cells (Fig. 3a). While we could not rule out the effect we observed could be due, at least in part, to a general effect of butyrate on gene expression, we did observe a sequence-dependent effect of butyrate on expression. Cells transduced with a FLAG-tagged version of “SCN1A” or the related voltage-gated sodium channel, similarly-sized transgene, *SCN5A*^29^, showed differential responses to butyrate treatment, with the former surpassing the latter in percent of cells expressing the reporter only after butyrate treatment, although we observed similar increases in MFI (Fig. 3b, c). As the flanking sequence elements of both LVVs were the same, including the promoter and emGFP reporter, we posit that the *SCN1A* sequence itself affects expression gene efficiency. This could in part be due to *SCN1A* sequence-specific elements that are associated with epigenetic repression or regulation.

**Figure 3:**
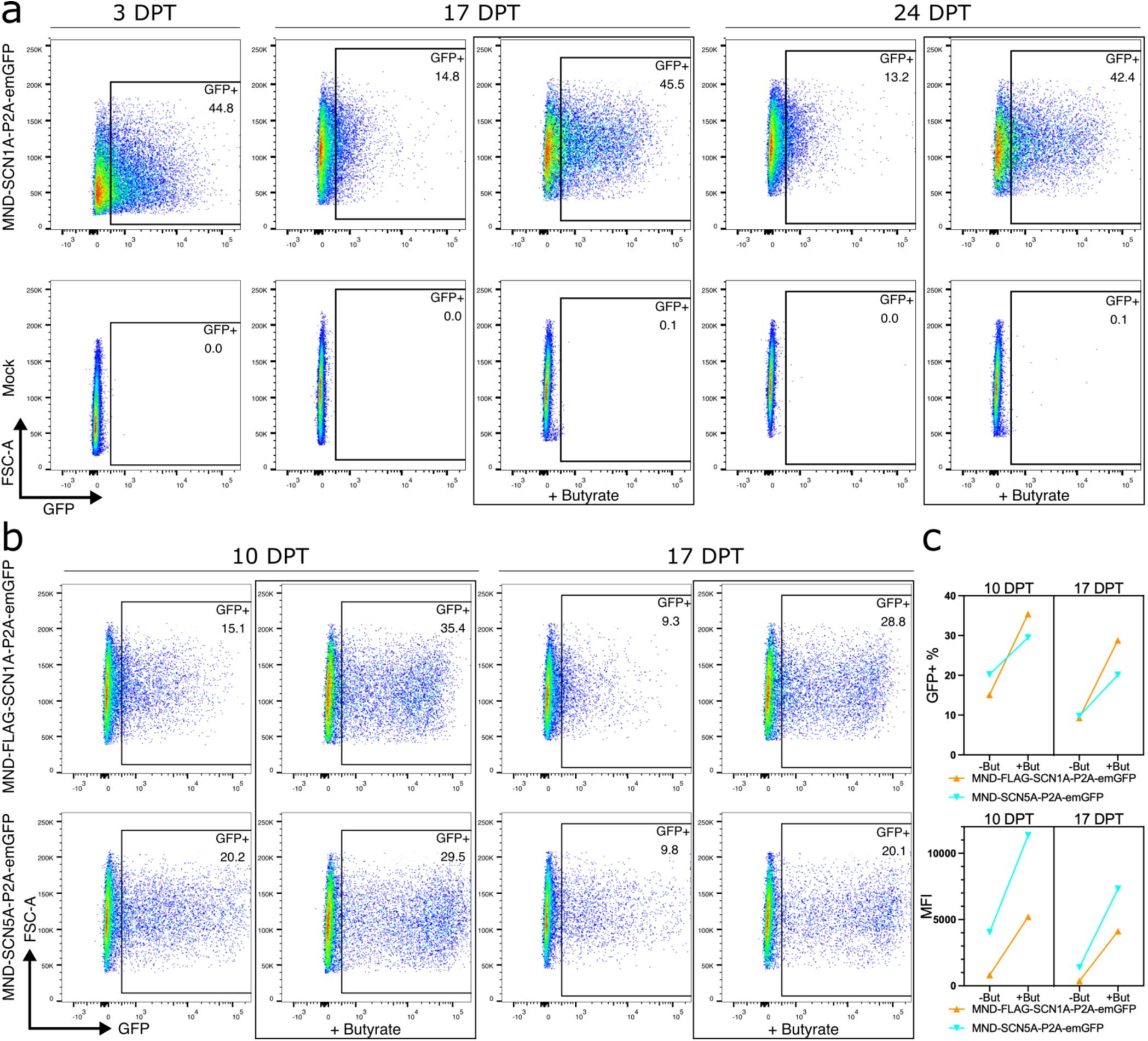
Butyrate helps counter the decay in expression of sodium channel constructs in a sequence-dependent fashion. (a) 5 mM sodium butyrate or no treatment was administered 24 hours before flow cytometry and GFP expression measured at the indicated days post-transduction (DPT) in 293T cells transduced with either MND-SCN1A-P2A-emGFP or mock. (b) GFP expression of 293T cells transduced with either MND-FLAG-SCN1A-P2A-emGFP or MND-SCN5A-P2A-emGFP at the indicated DPT, with or without 5 mM sodium butyrate administered 24 hours before flow cytometry. (c) Summary of data in (b), namely percentage of GFP+ and mean fluorescence intensity (MFI) of transduced cells.

### Adding a ubiquitous chromatin opening element upstream of the LVV MND promoter results in a trend of higher emGFP reporter expression and more responsiveness to butyrate treatment

Hypothesizing that sequence-dependent elements could contribute to epigenetic repression of our “SCN1A” sequence, we introduced a ubiquitous chromatin opening element (UCOE)^20^ immediately upstream of the MND promoter in our transgene cassette. UCOEs have been shown to help proximal elements, such as a promoter, resist DNA methylation, thus enhancing expression^30^. While we did not observe any significant increase in emGFP reporter expression in LVV-transduced cells due to the UCOE, there was a trend of higher reporter expression over time, compared to the original, non-UCOE “SCN1A” LVV, which was accentuated with butyrate treatment (Fig. 4). These data suggest that, while the UCOE may help boost expression in the context of LVVs, its potential to enhance expression may be limited depending on the context of the downstream transgene sequence, such as repressive elements present within the *SCN1A* sequence itself, which could lie outside the range of or counteract the beneficial effects of the UCOE.

**Figure 4:**
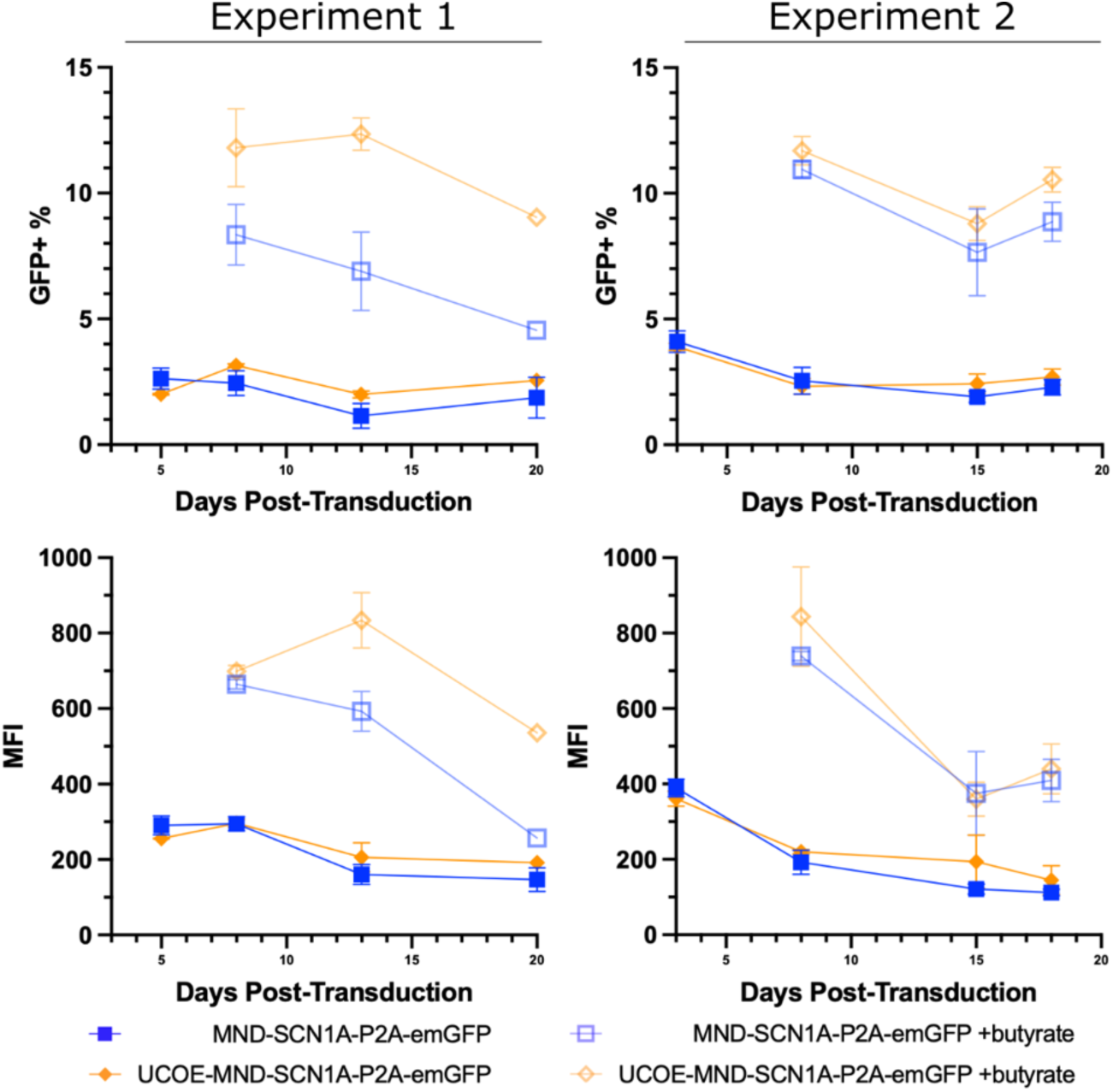
Adding a UCOE element upstream of promoter results in trends of higher reporter expression and more responsiveness to butyrate treatment. Percent GFP+ (top) and mean fluorescence intensity (MFI, bottom) of 293T cells transduced with either MND-SCN1A-P2A-emGFP or UCOE-MND-SCN1A-P2A-emGFP at the indicated days post-transduction, with or without 5 mM sodium butyrate administered 24 hours before flow cytometry. Means shown with standard deviation (*n* = 2 well replicates where error bars present).

## Discussion

This study demonstrates that *SCN1A* transgenes can be delivered to human cells via lentiviral vector (LVV)-mediated transduction for expression of full-length Na_V_1.1 protein that traffics to the membrane and results in electrophysiologic changes consistent with ion channel expression. We observed that the sequences of the channel proteins we tested, despite identical flanking elements in the LVV transgene cassette, impacted the efficiency of expression in transduced cells. Moreover, expression declined substantially over time in culture regardless of the exact composition of the *SCN1A* transgene sequence.

It is possible that expression in a more physiologically relevant context may help sustain expression of *SCN1A* transgenes. Expression of sodium channel β subunits^31^, or expression in a neuronal cell line may alleviate *SCN1A* transgene expression problems found in 293T. However, anecdotally, we were unable to express *SCN1A* LVVs in the Neuro-2a (murine neuroblast) cell line to levels comparable to those in 293T cells. Some preliminary analysis also suggested co-expression of *SCN1B* and *SCN2B* with *SCN1A* did not result in higher Na_V_1.1 protein (compared to *SCN1A* expression alone). This again reinforces our proposal that the regulation of *SCN1A* transgene expression may be primarily sequence-dependent.

While we found splicing of the LVV genome itself (i.e. in its DNA form post-transduction in 293T cells), it remains possible that splicing at the mRNA level (i.e., transgene expression) could be contributing to the low observed expression in transduced 293T cells. However, again anecdotally, we did not detect aberrant mRNA splice products in the few predicted alternative splice sites we amplified that were flagged as possible splice sites by a webtool^32^. Even if splicing at the mRNA level is contributing to lower overall expression of the *SCN1A* transgene, this would not explain why expression decreases over time.

A question we could not fully address was whether the lower *SCN1A* transgene expression we observed in our original “*SCN1A*” construct compared to the optimized (“Opt”) construct was due to lower transgene transcription in transduced cells, lower protein translation, or a combination of the two. While our results with butyrate treatment may imply a sequence-dependent effect on transcription, it is also possible that features of the mRNA itself, including codon optimization^33^, length^34^, secondary structure^35^, location of a synthetic intron^36^, other cryptic regulatory elements, etc. could impact translation.

While we suspect that our findings are not unique to LVVs, other studies have found some success delivering *SCN1A* via other vectors, including adenoviruses and adeno-associated virus (AAV). Two studies using adenovirus vector expressing *SCN1A* under the control of neuron-specific promoters each showed some restoration of Dravet syndrome (DS) phenotype in mouse models, when administered intracranially^37,38^. However, a major limitation of adenoviral vectors is that their safety profile, especially for CNS indication, has not been established in clinical trials. A recent study employing a split-intein system to express *SCN1A* in two AAV vectors, due to the lower genetic capacity of AAVs^39^, also showed improvement of DS phenotypes in a mouse model^24^. What was not documented in these studies with other vectors was whether *SCN1A* expression diminished over time.

The promise of LVVs for gene therapy comprises three key features: their relatively large payload, stable expression, and targetability. While the first of these features enabled this study, namely, delivery of functional *SCN1A* in a single delivery platform to cells, there remains questions as to why LVV *SCN1A* expression did not achieve stable expression over time. If additional progress can be made towards overcoming this hurdle, this would enable future studies that address the third feature—targeting the vector to specific neuronal cell types that would especially benefit from therapeutic *SCN1A* transgene expression. Our findings highlight a new potential approach for treating Dravet syndrome and the areas where additional studies are needed to achieve new milestones in the quest for cures for *SCN1A* deficiency and related channelopathies.

---

**Supplementary Table S1.**
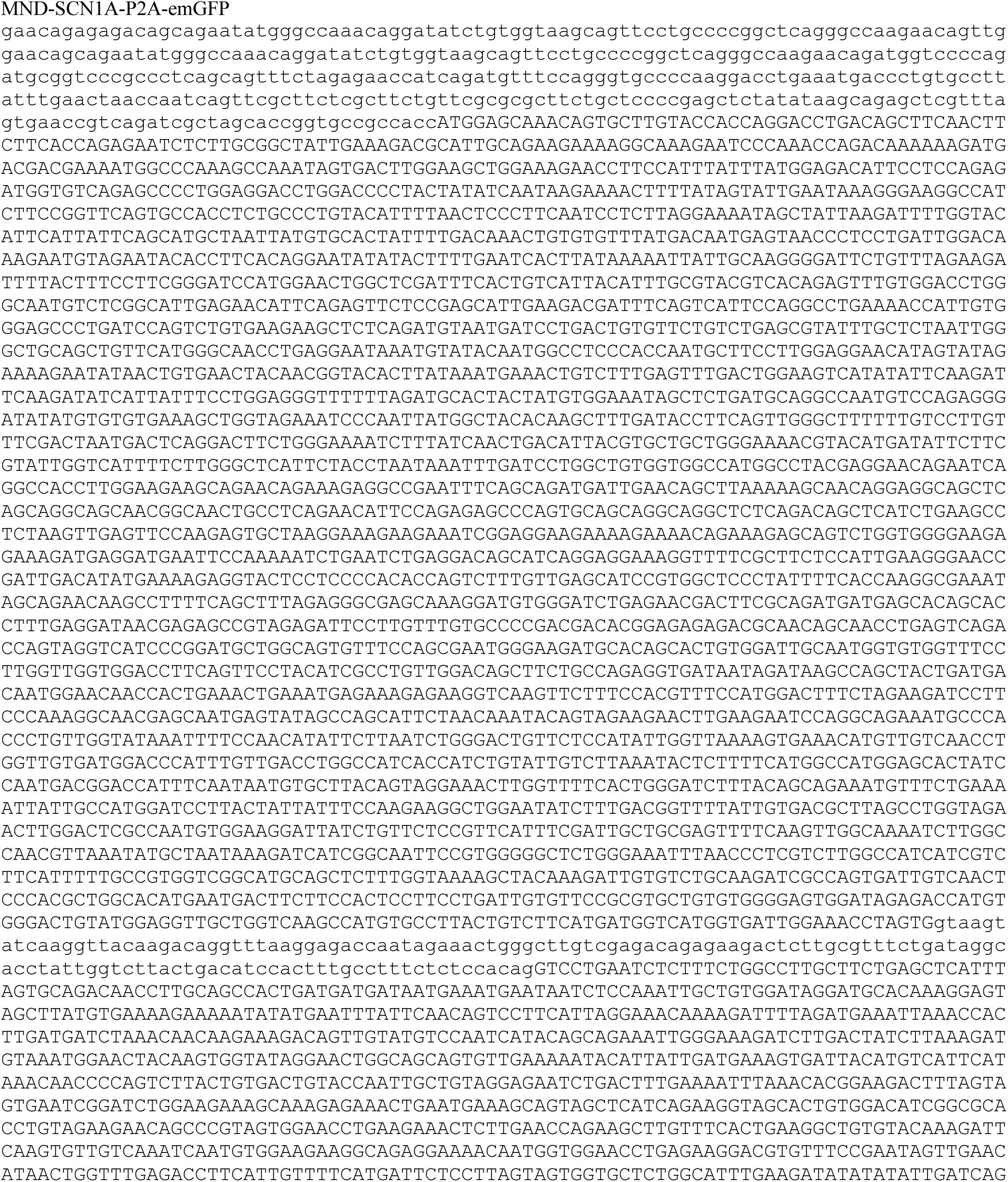
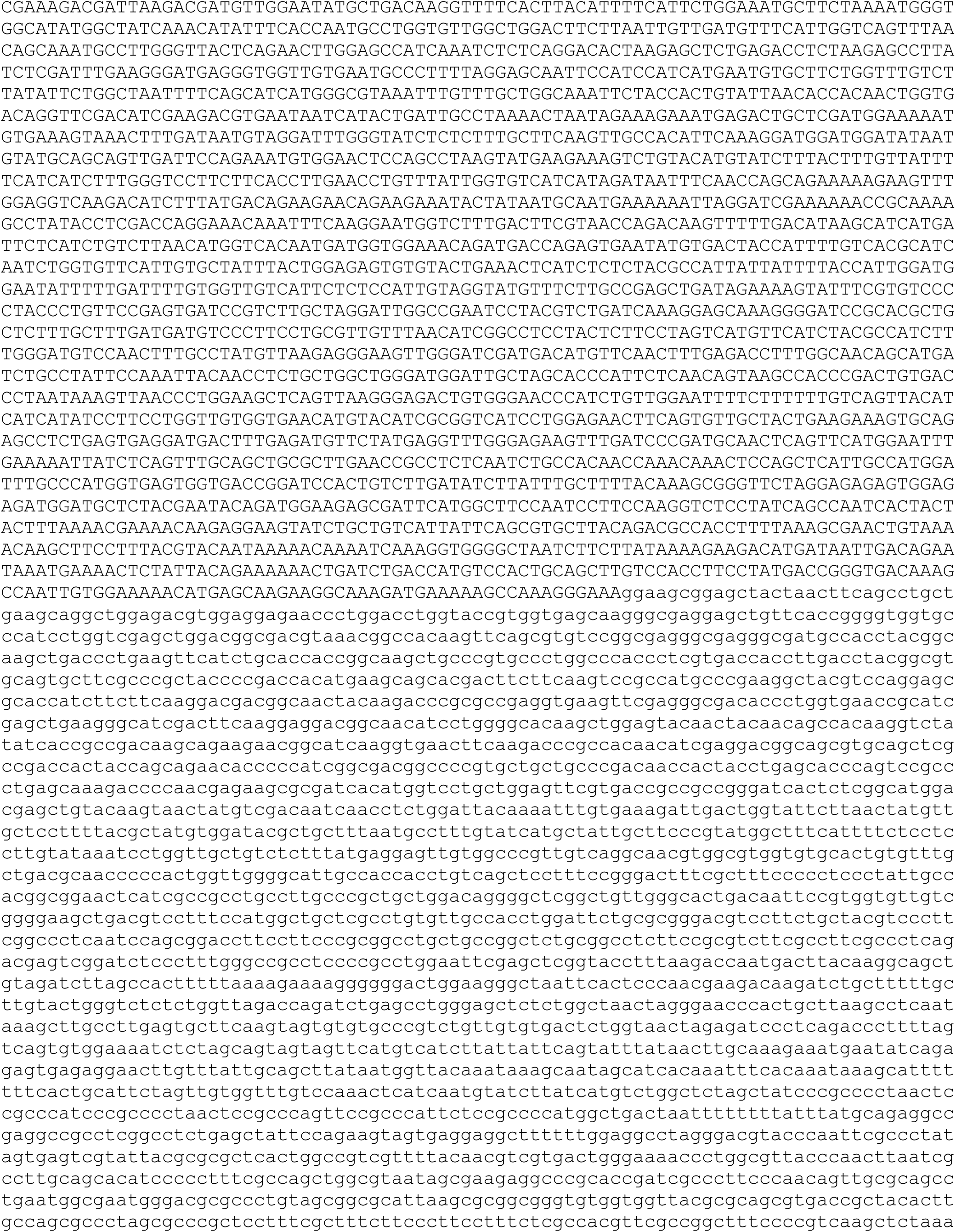
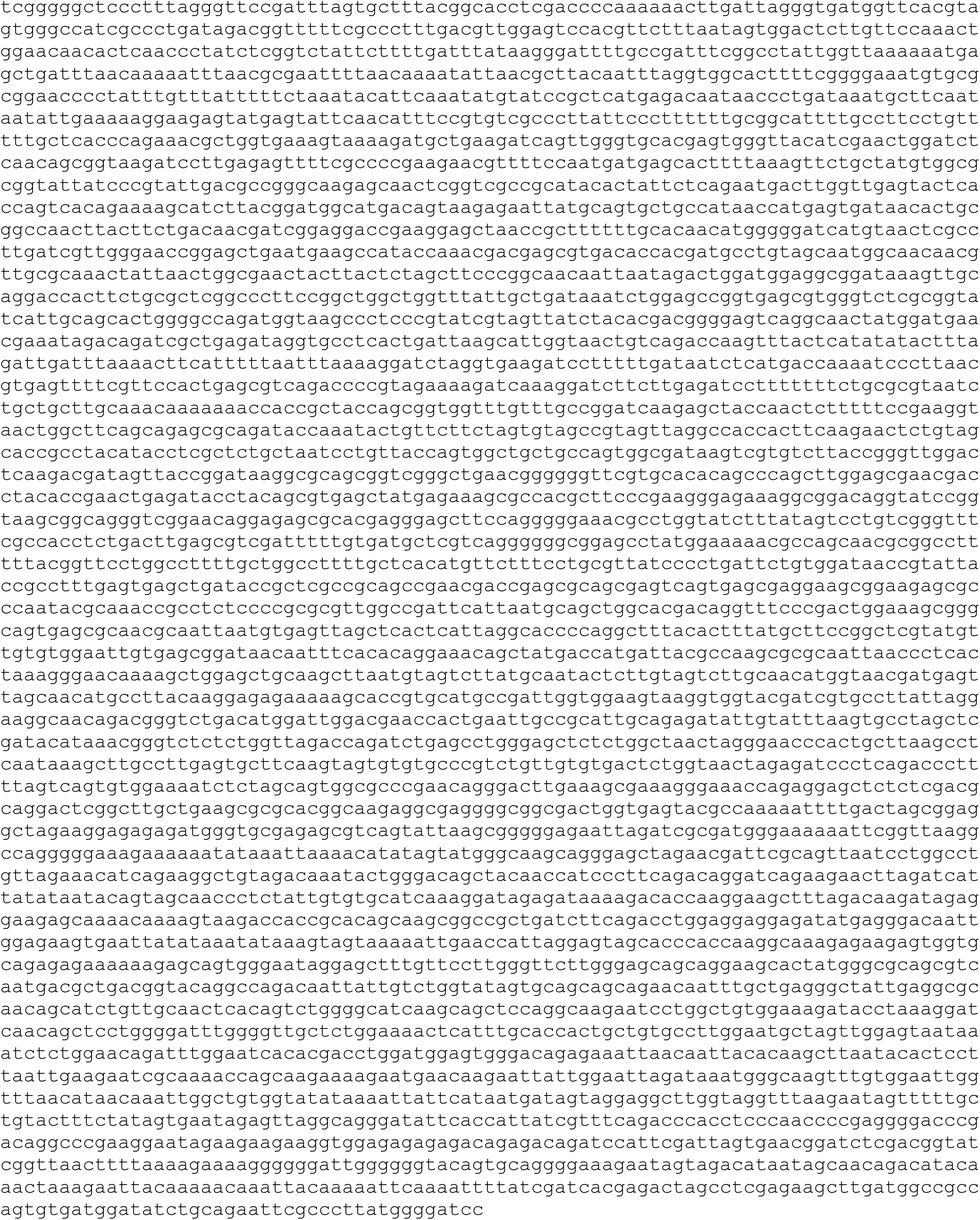
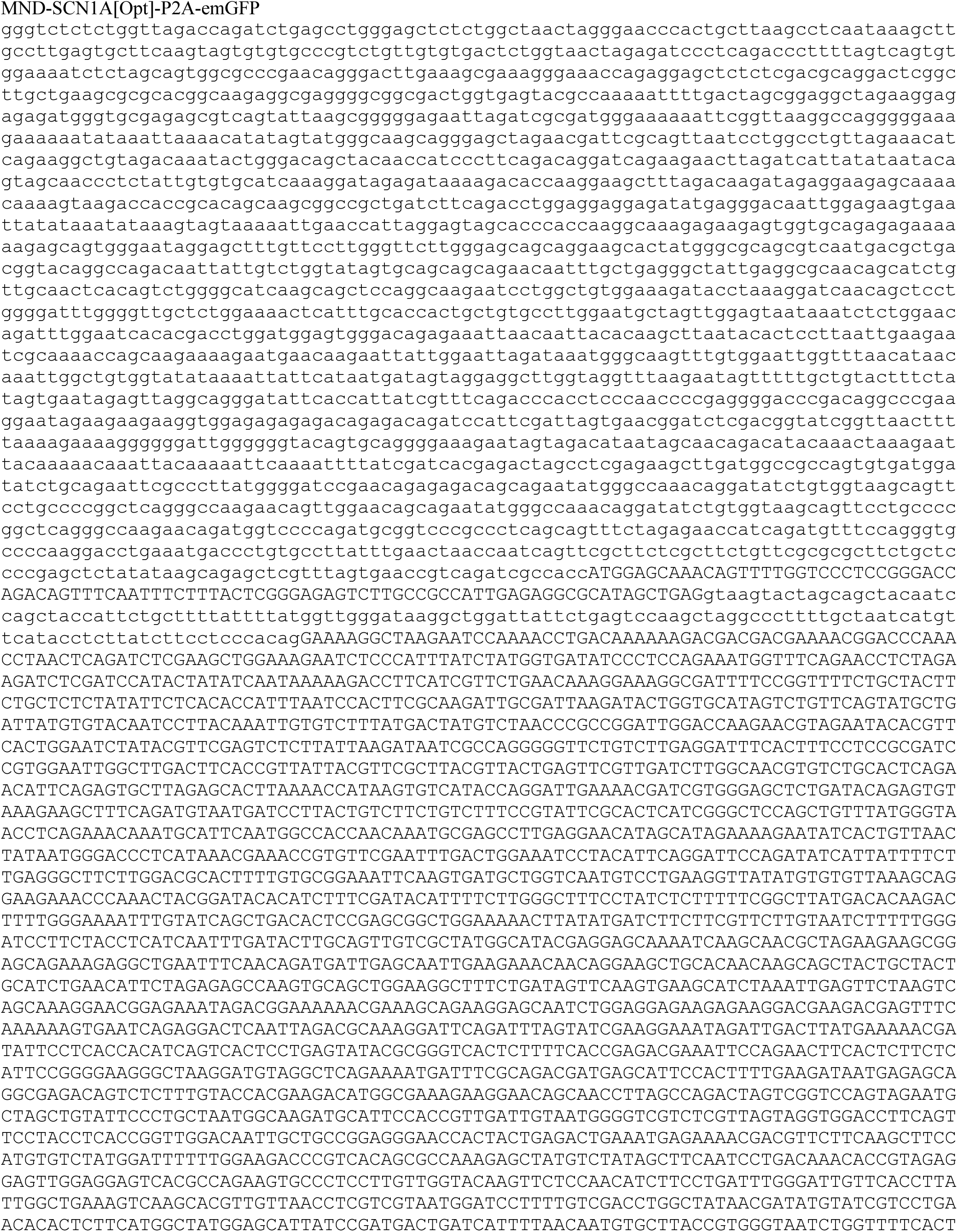
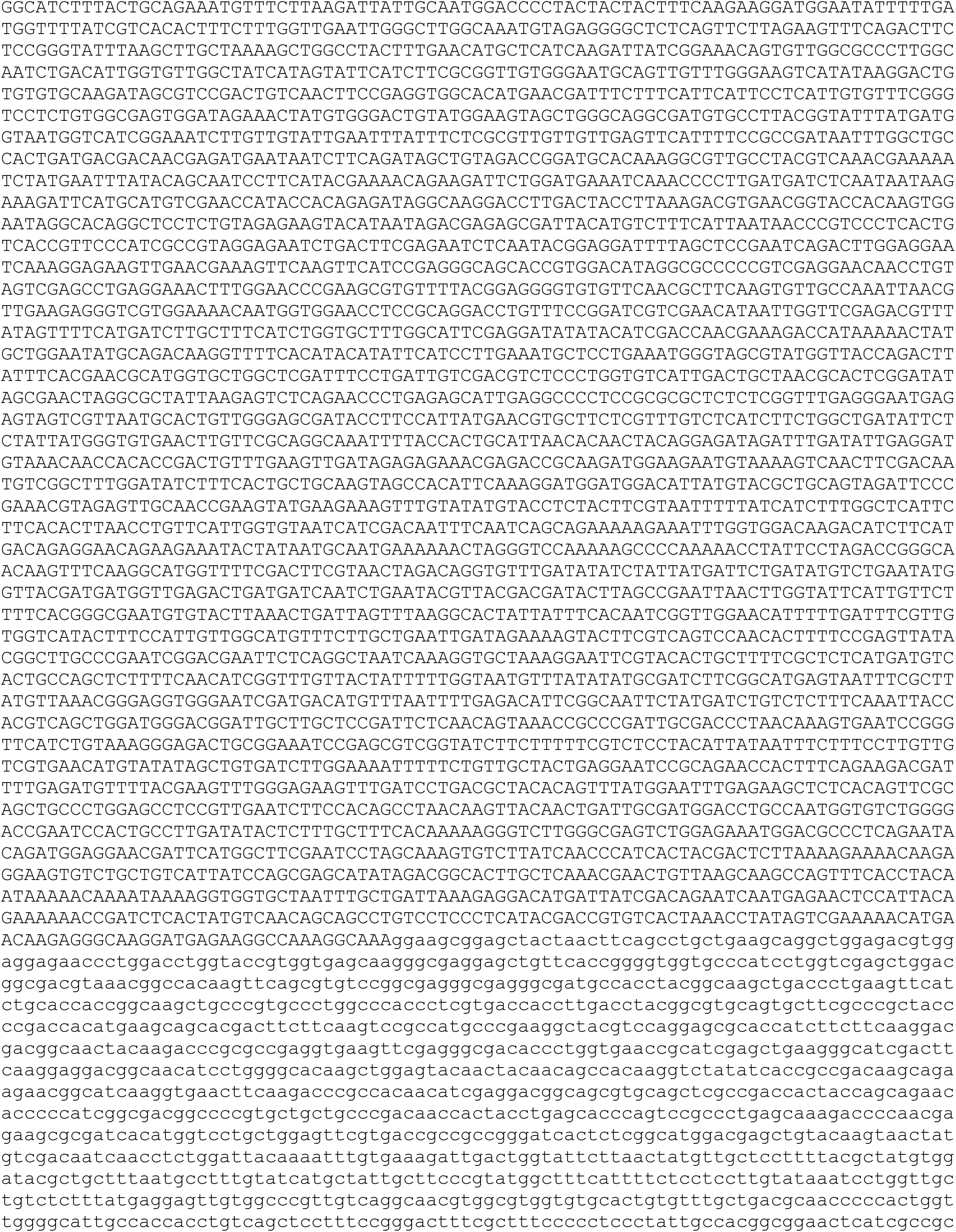
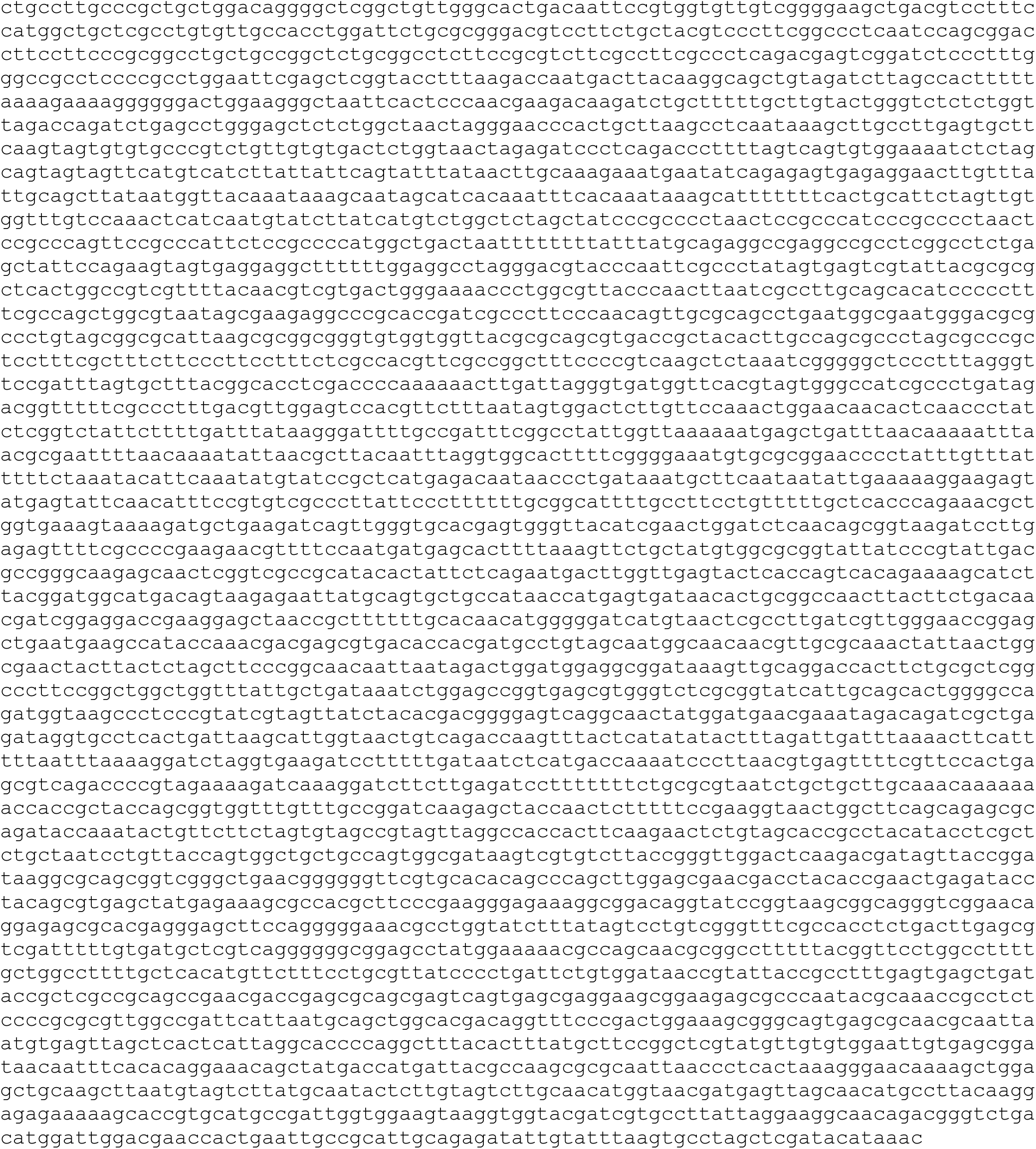
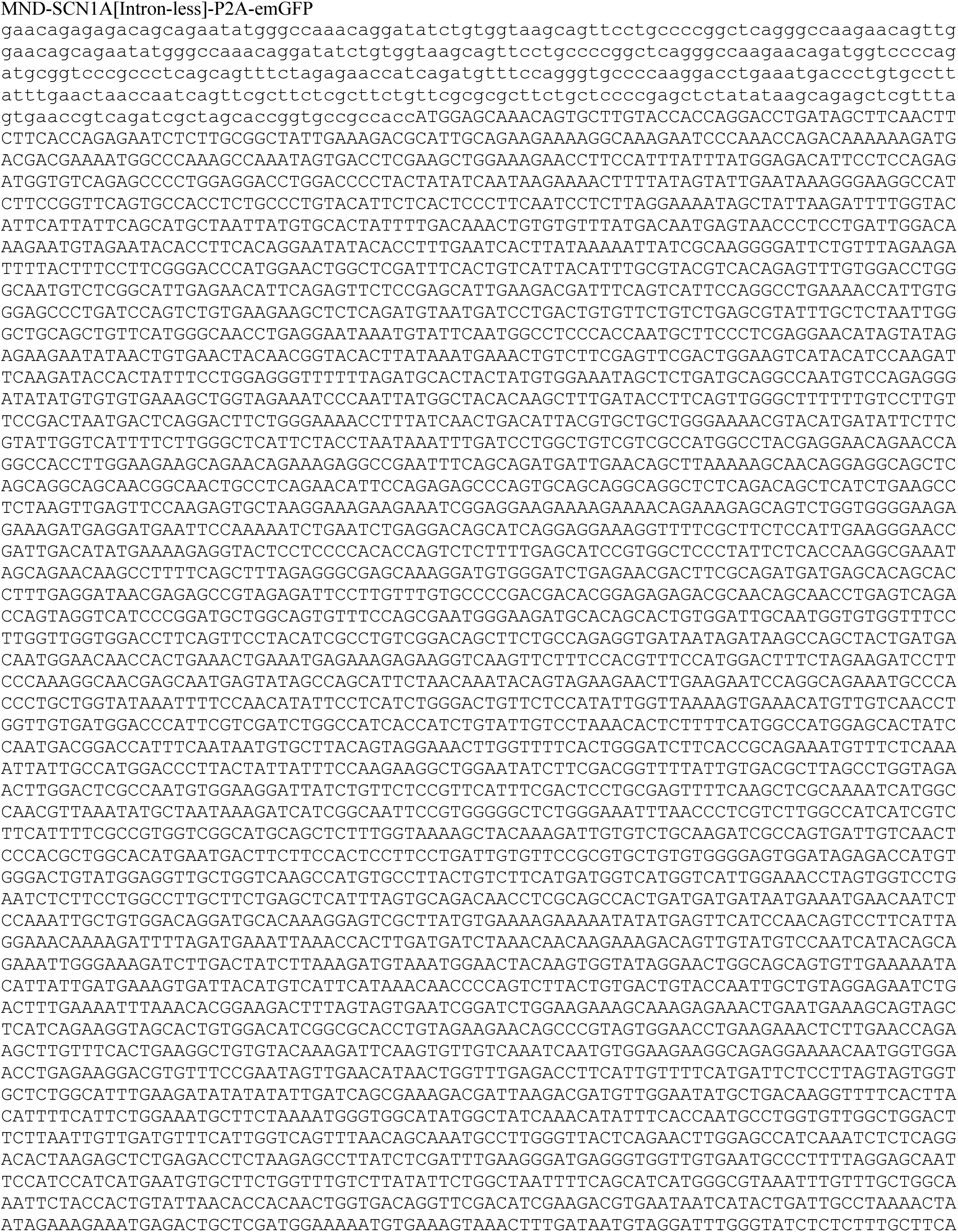
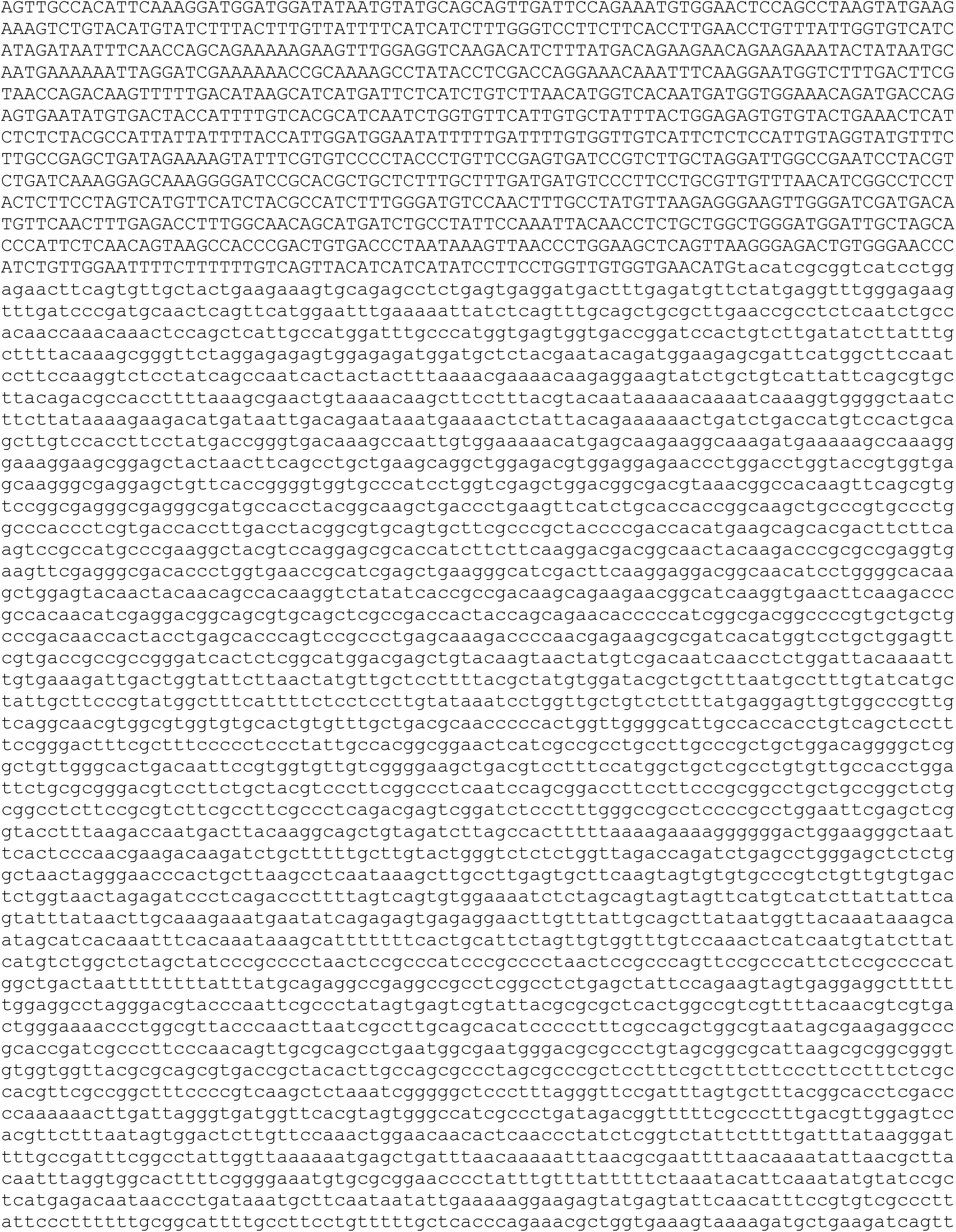
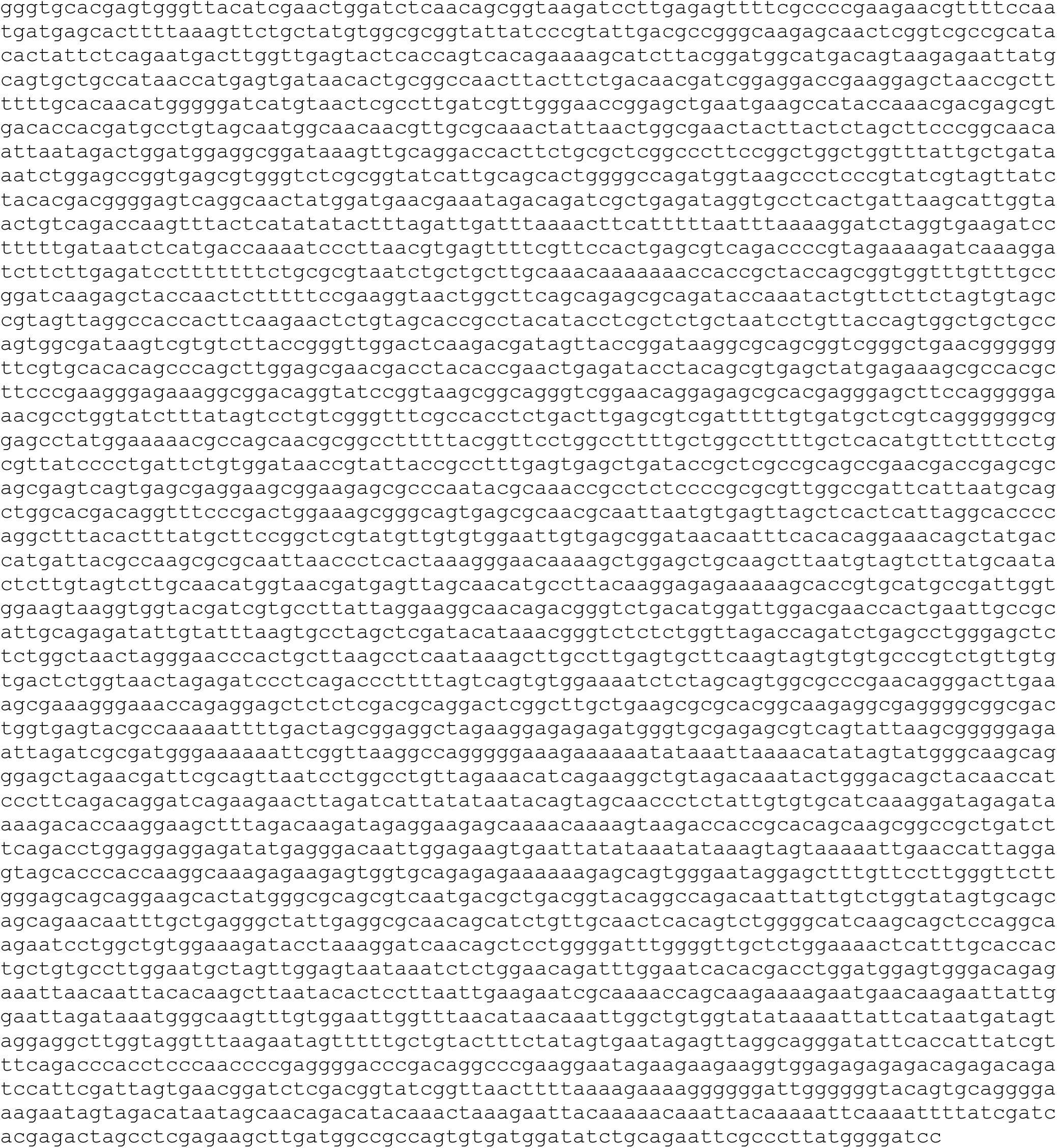
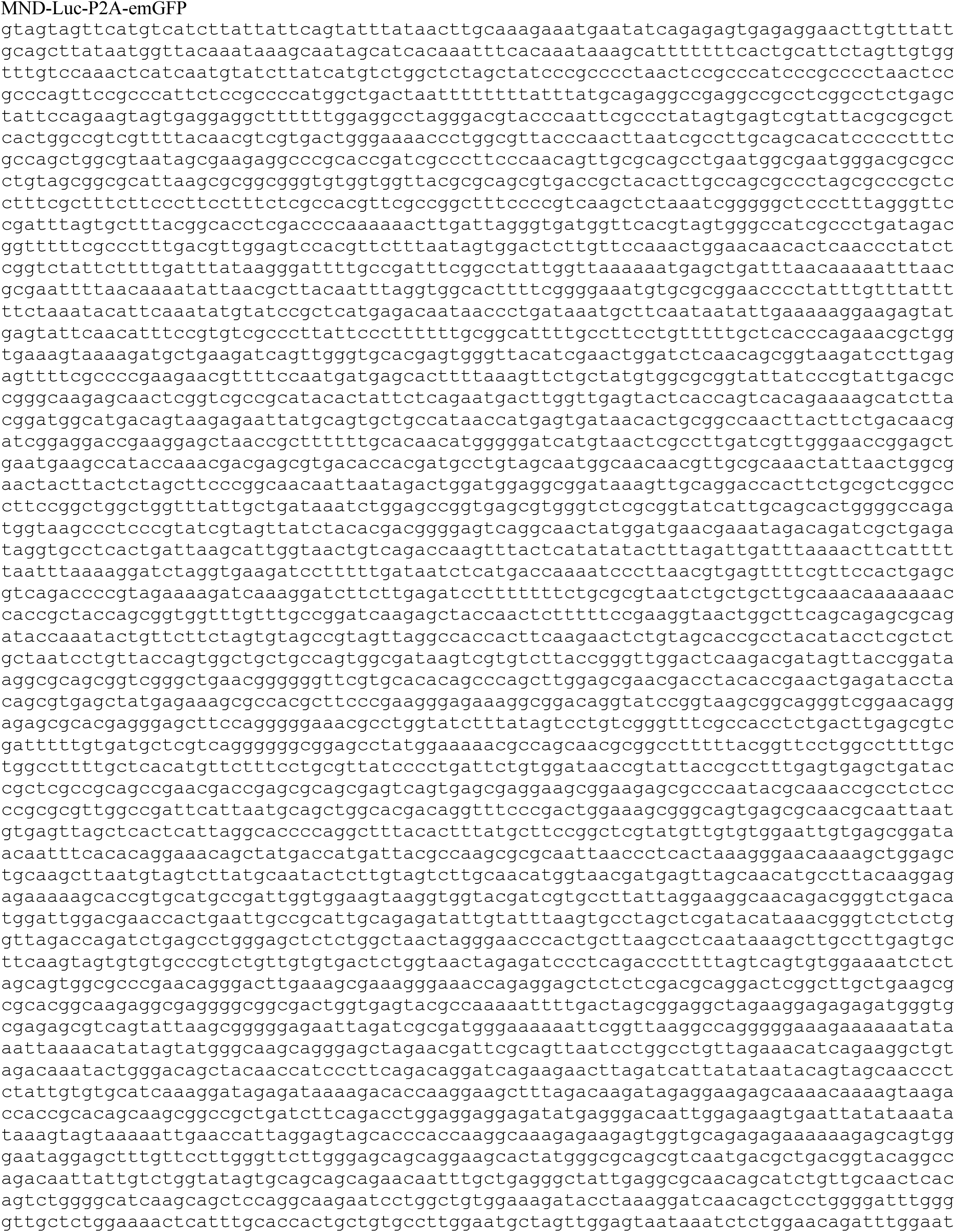
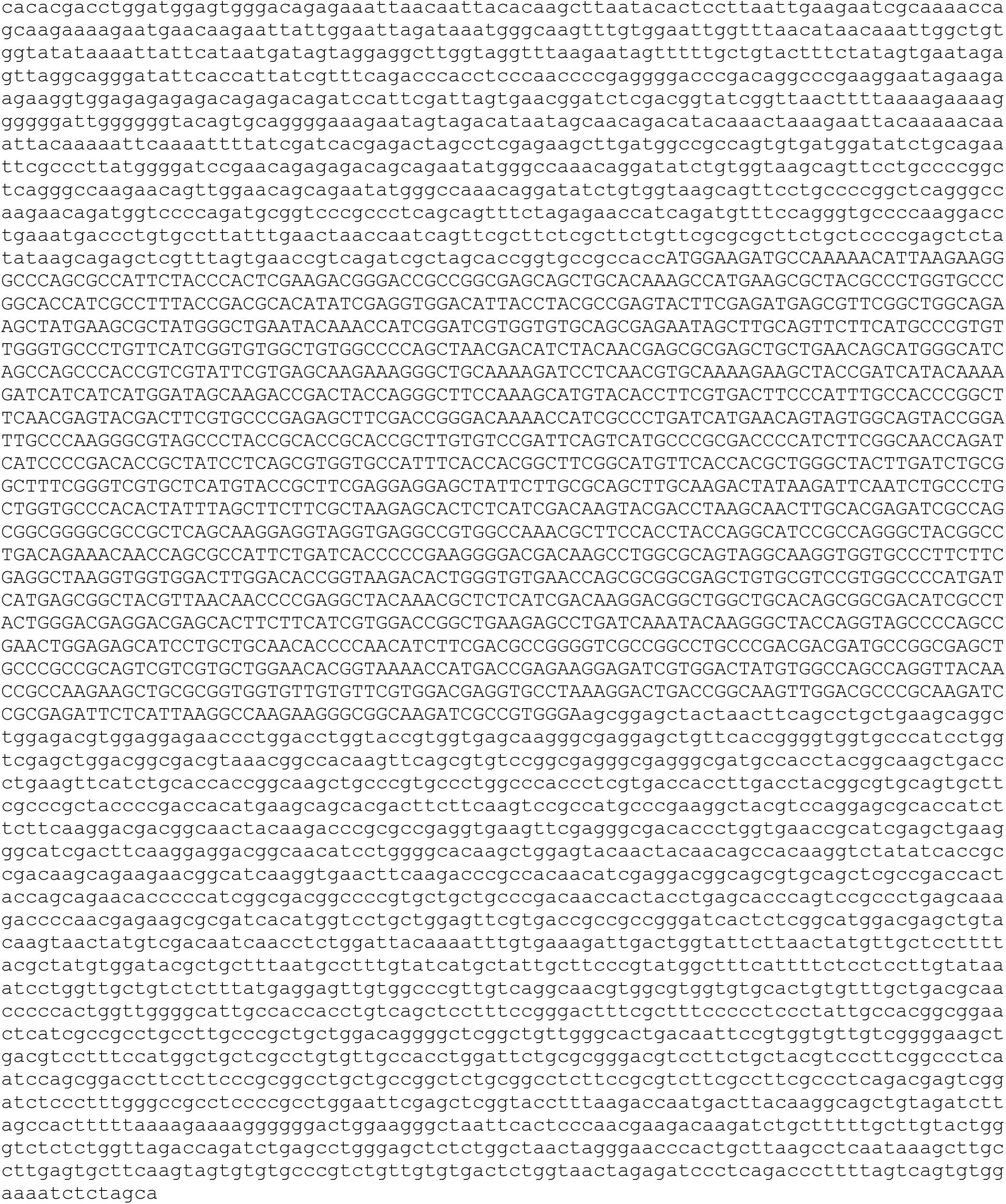
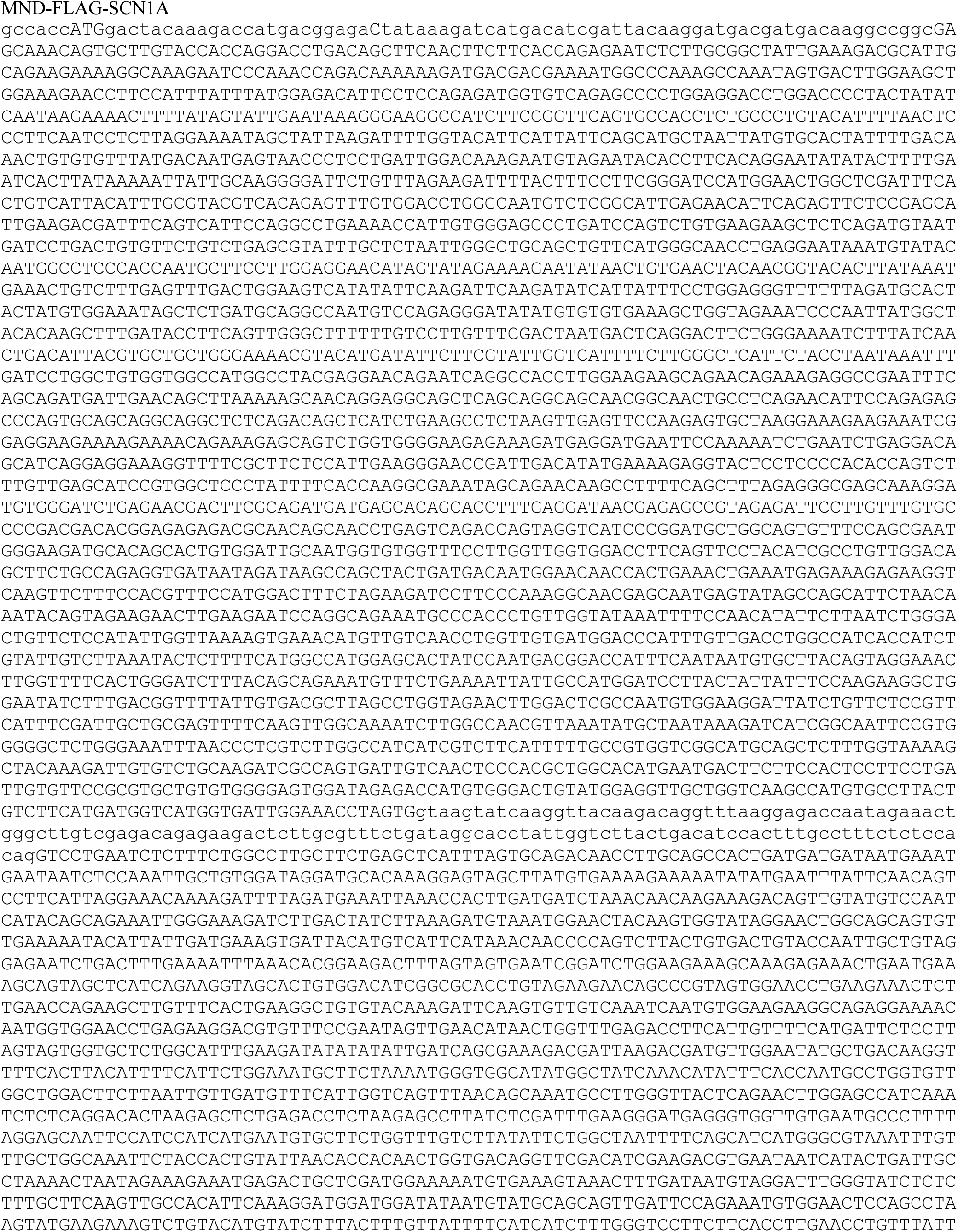
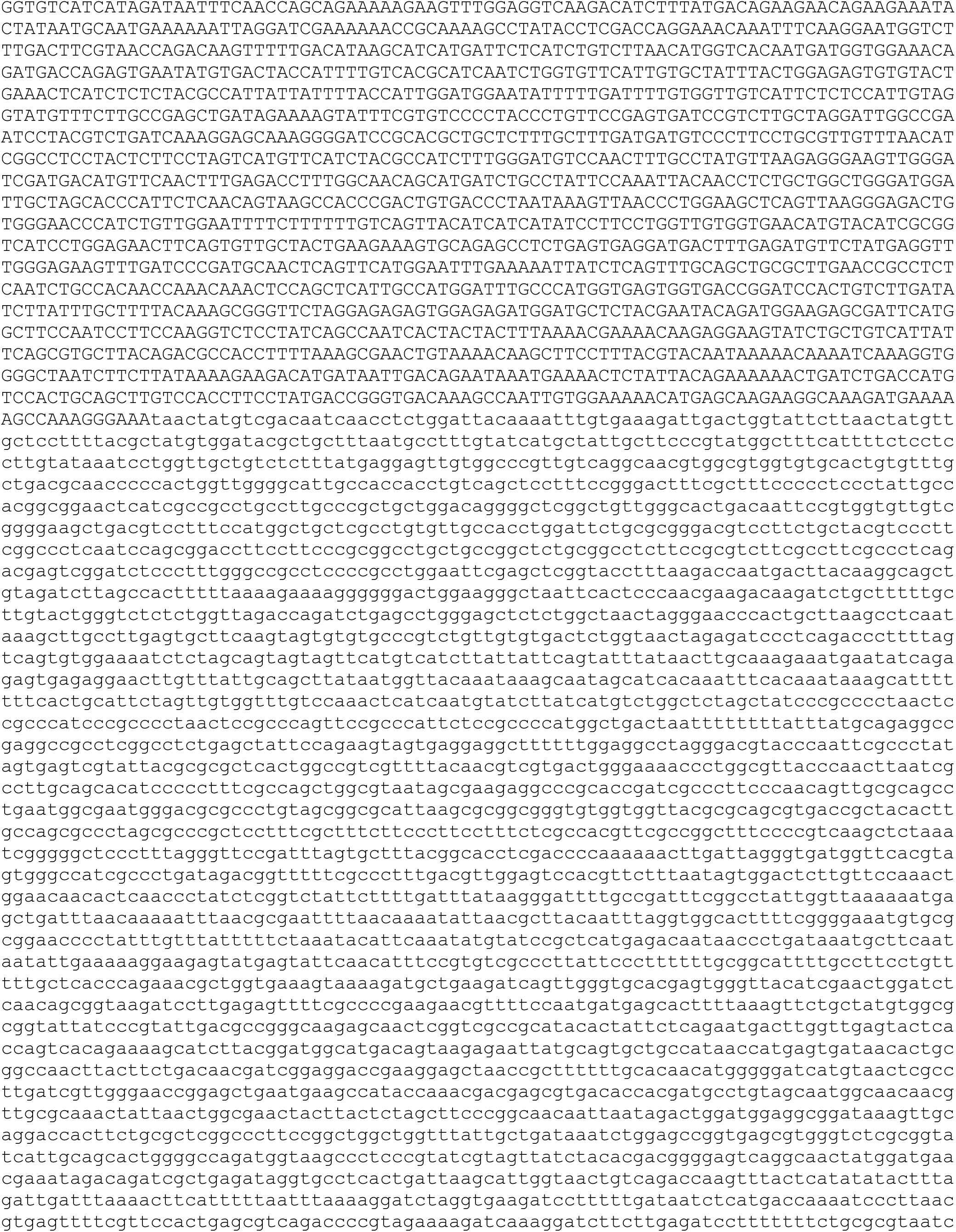
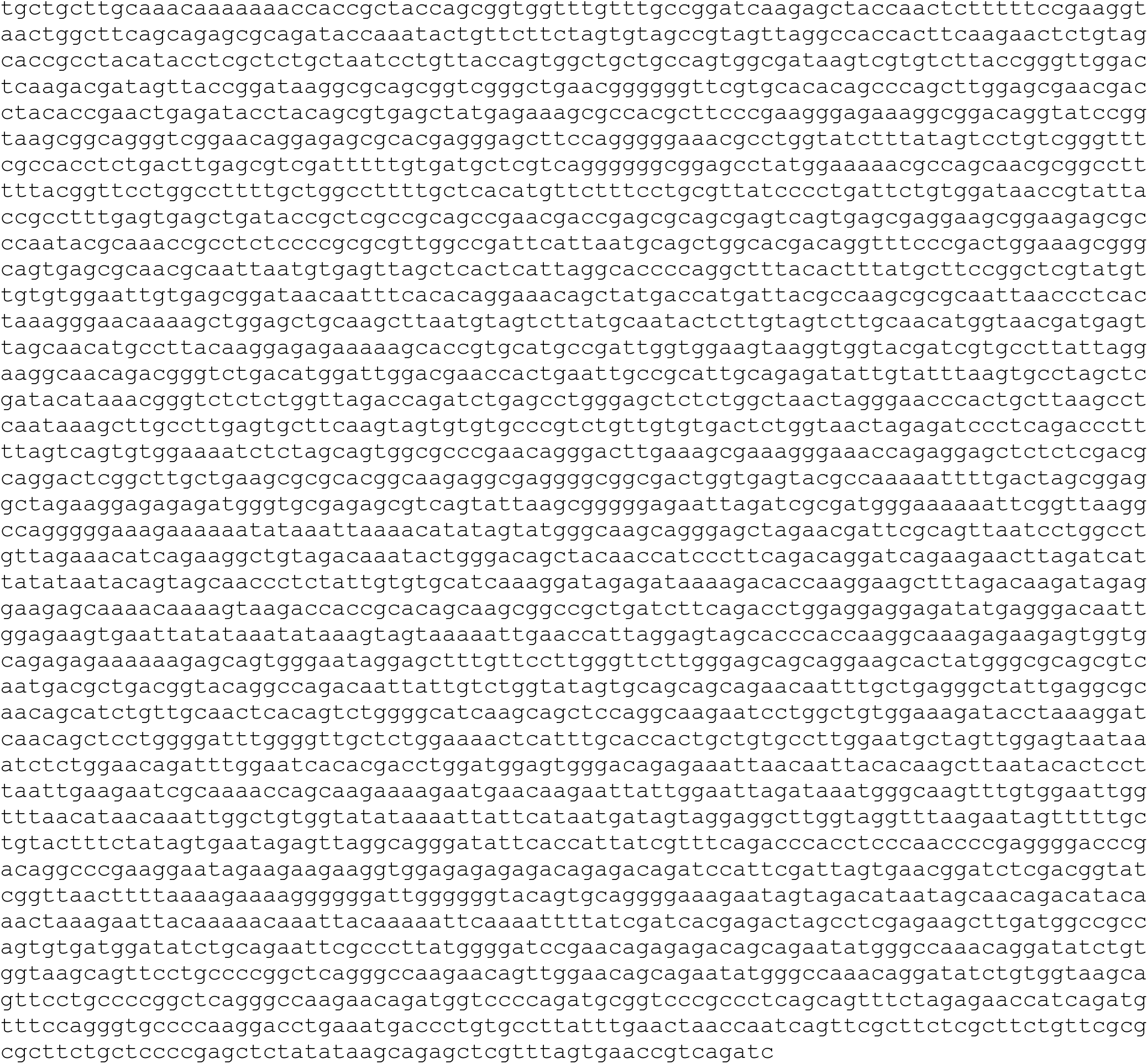
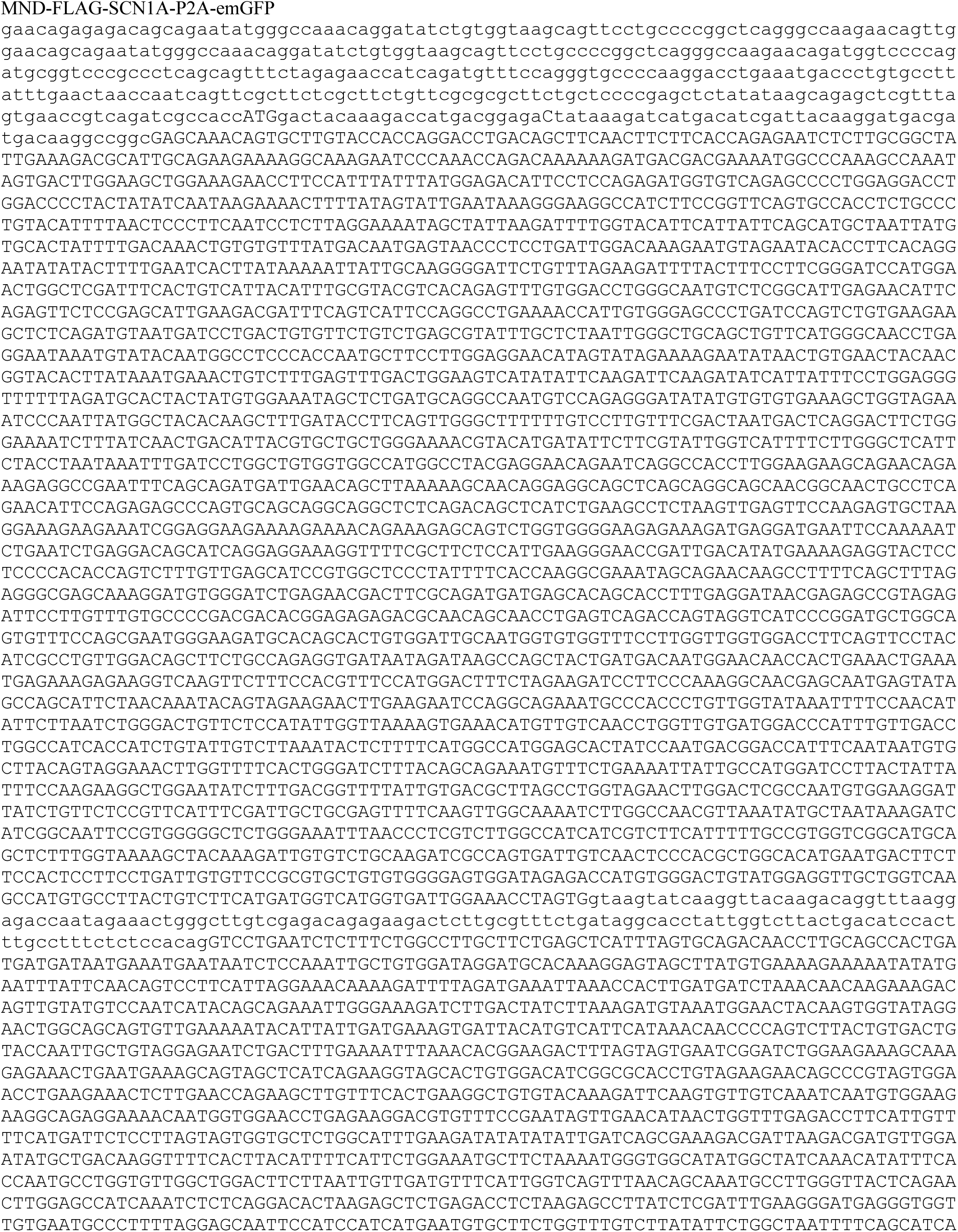
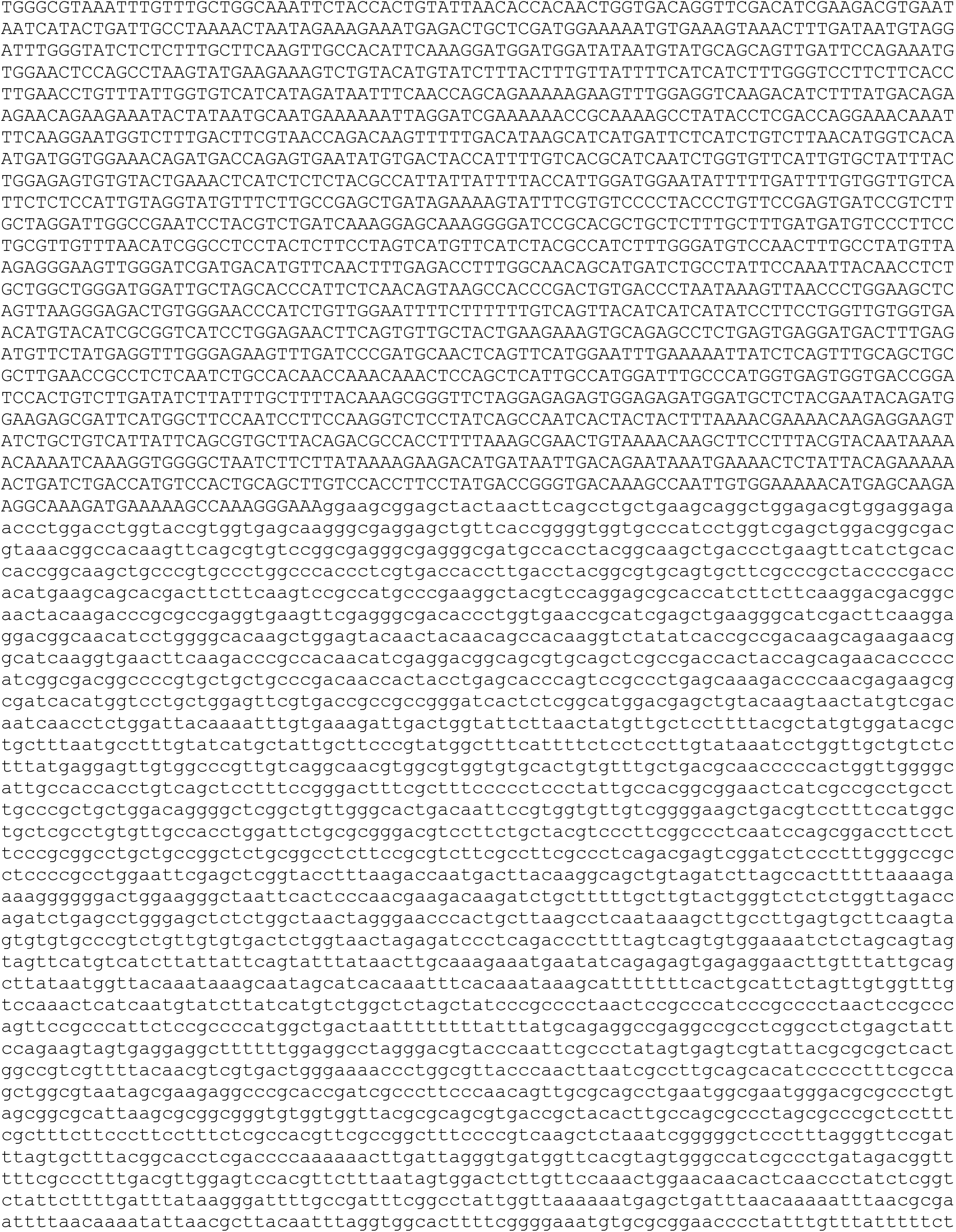
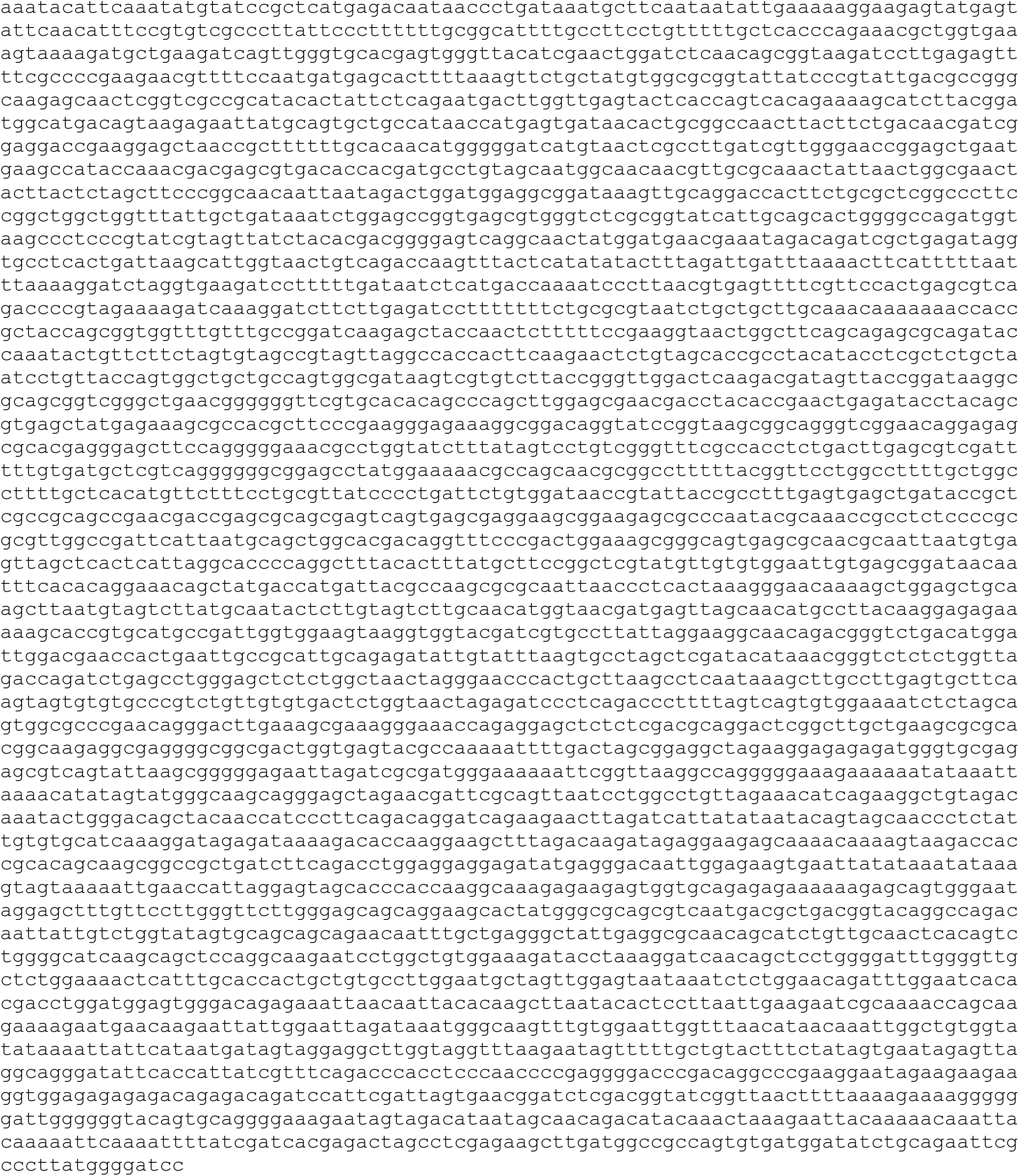
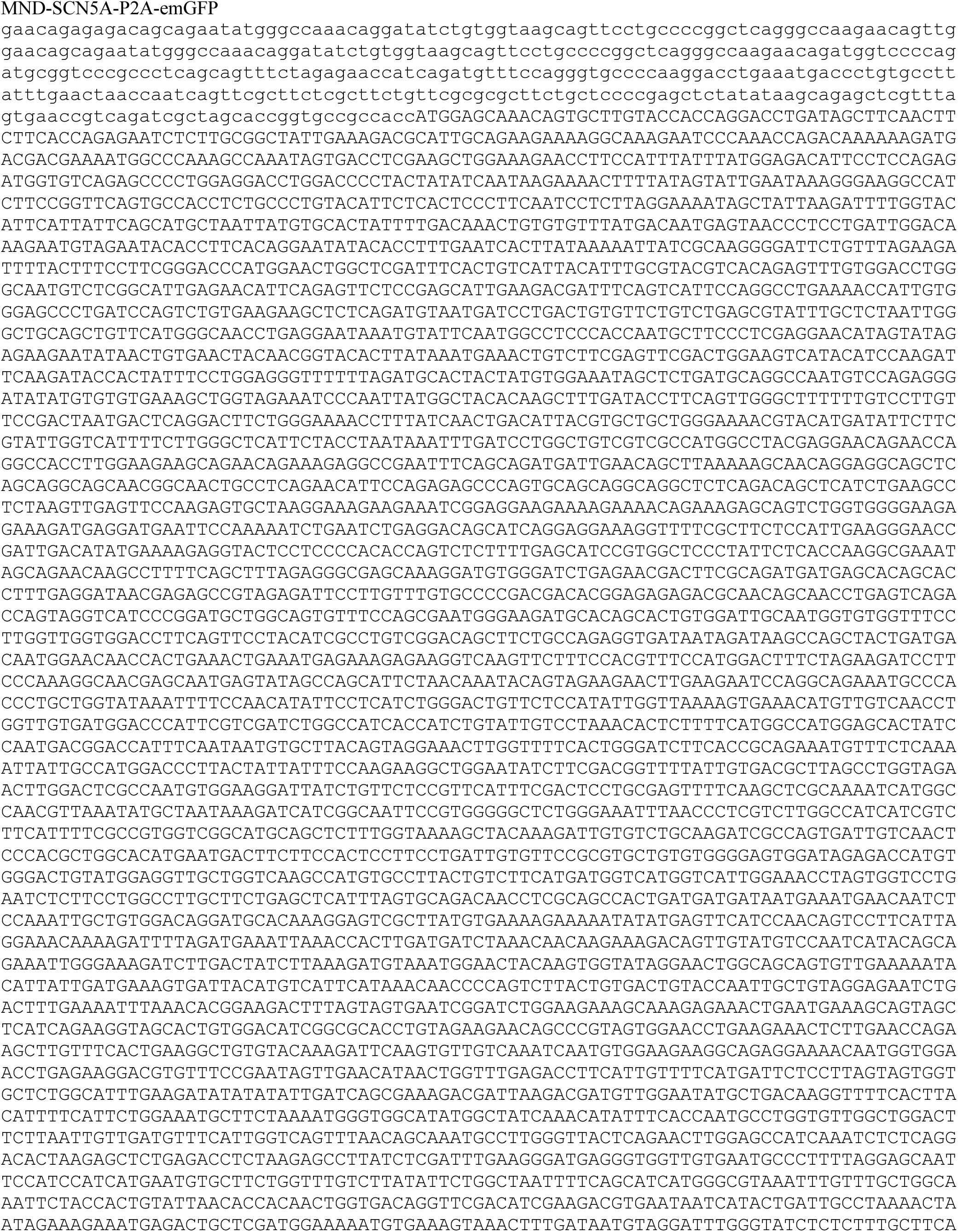
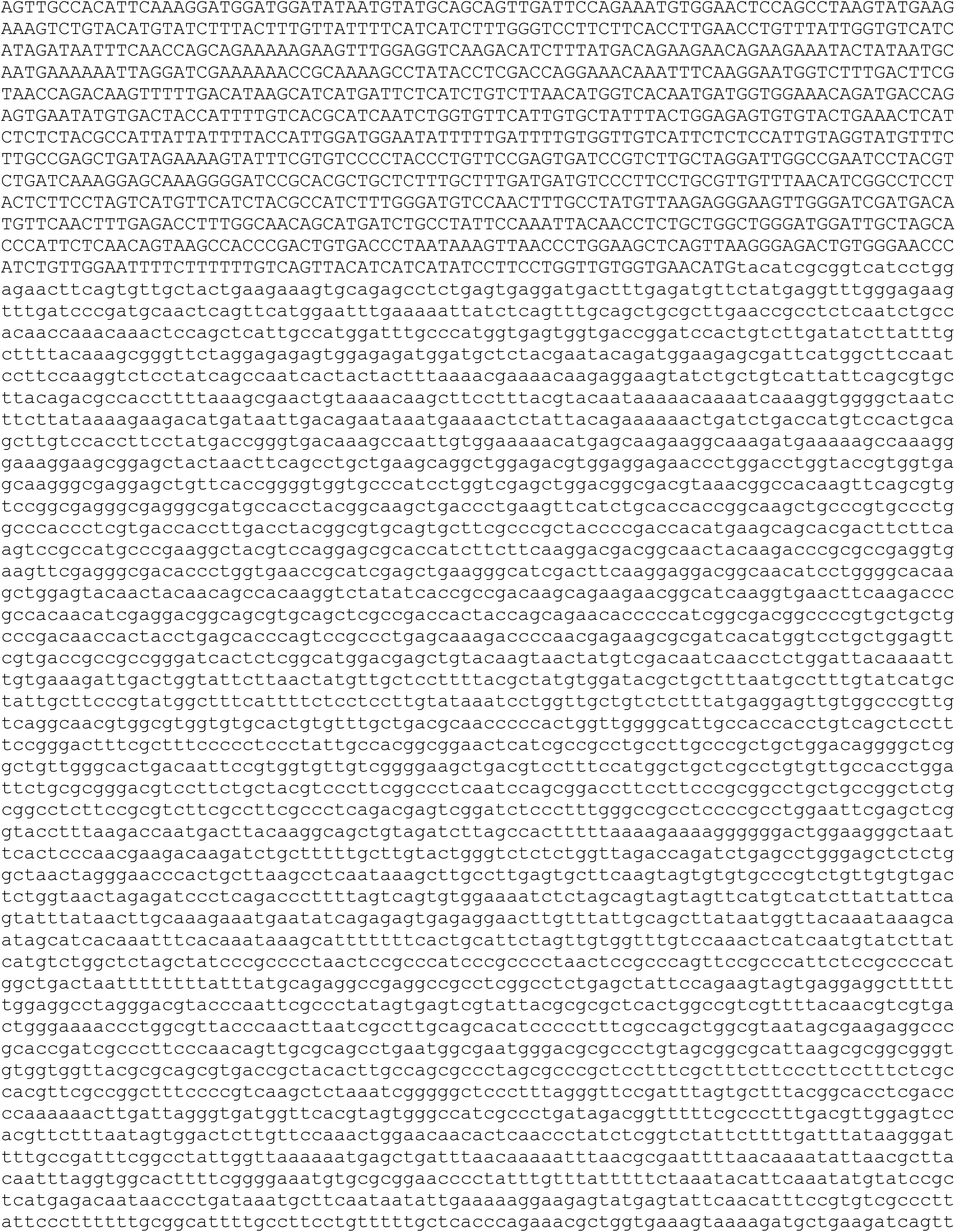
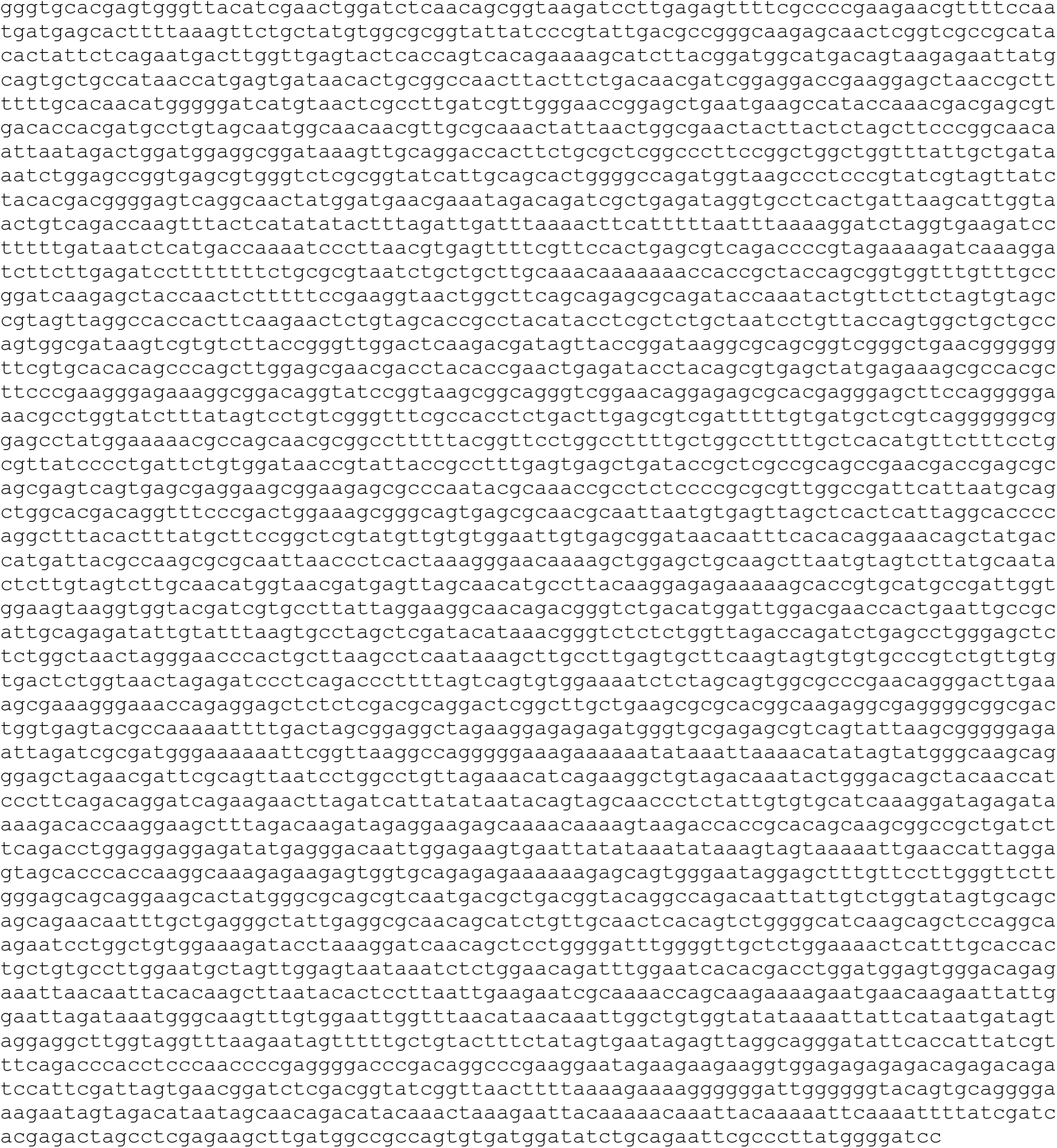
Sequences of *SCN1A* plasmid constructs used in this study. Note that the *SCN1A* or *SCN5A* (or luciferase for the control) coding sequence is denoted in uppercase. Note that there may have been backbone mutations outside of the region spanning the lentiviral transgene cassette.

## Notes

### Competing Interest Statement

The authors have declared no competing interest.

### Summary of Updates

Edited title (again) to remove italics on the wrong word.

